# The mouse dorsal peduncular cortex encodes fear memory

**DOI:** 10.1101/2023.07.24.550408

**Authors:** Rodrigo Campos Cardoso, Zephyr R. Desa, Brianna L. Fitzgerald, Alana Moore, Jace Duhon, Victoria A. Landar, Roger L. Clem, Kirstie A. Cummings

## Abstract

The rodent medial prefrontal cortex (mPFC) is a locus for both the promotion and suppression (e.g. extinction) of fear and is composed of four anatomically distinct subregions, including anterior cingulate 1 (Cg1), prelimbic (PL), infralimbic (IL), and the dorsal peduncular (DP) cortex. A vast majority of studies have focused on Cg1, PL, and IL. The Cg1 and PL have been implicated in the promotion of fear, while the IL has been linked to a role in the suppression, or extinction, of fear. Due to its anatomical location ventral to IL, the DP has been hypothesized to function as a fear-suppressing brain region however, no studies have explicitly tested its role in this function or in the regulation of memory generally. Moreover, some studies have pointed towards a dichotomous role for ventral mPFC in the dual suppression and promotion of fear, but the mechanisms underlying these opposing observations remains unclear. Here, we provide evidence that the DP paradoxically functions as a cued fear-encoding brain region and plays little to no role in fear memory extinction. By using a combination of cFos immunohistochemistry, whole-cell brain slice electrophysiology, fiber photometry, and activity-dependent neural tagging, we demonstrate that DP neurons exhibit learning-related plasticity, acquire cue-associated activity across learning and memory retrieval, and that DP neurons activated by learning are preferentially reactivated upon fear memory retrieval. Further, optogenetic activation and silencing of fear learning-related DP neural ensembles drives the promotion and suppression of freezing, respectively. Overall, these data suggest that the DP plays an unexpected role in fear memory encoding. More broadly, our results reveal new principles of organization across the dorsoventral axis of the mPFC.

## Introduction

Several studies have pointed towards the rodent medial prefrontal cortex (mPFC) as a locus important for both the promotion and suppression of fear. The mPFC is comprised of four anatomically and functionally distinct subregions including the anterior cingulate 1 (Cg1), prelimbic cortex (PL), infralimbic cortex (IL), and the dorsal peduncular cortex (DP). Numerous studies have largely ascribed the dichotomous role for the mPFC to PL and IL subdivisions, which play a role in the promotion and suppression of fear, respectively ^1–5^. Similar to the PL, Cg1 has also been implicated in signaling fear ^6,7^. In contrast, the specific role of the DP has largely been overlooked. The roles for Cg1 and PL in the promotion of fear and IL in the suppression of fear have led to the proposal that mPFC exhibits an organized functional architecture across the dorsoventral axis. Therefore, due to its anatomical location ventral to IL, some have hypothesized that DP also functions to suppress fear ^8^. Despite this, the role that DP might play in fear suppression or some other aspect of memory has largely not been tested (but see recent study^9^).

In contrast to its proposed role in safety learning, several studies suggest a role for the DP in processing aversive stimuli. Following social defeat stress, the DP, along with the region immediately ventral, the dorsal tenia tecta (DTT), have been implicated in driving physiological sympathetic responses, including thermoregulation and cardiac responses ^10–12^. Moreover, recent studies have also implicated the DP/DTT in driving anxiety-like behaviors^9^ and circuits of fight or flight^13^. Given the central role for autonomic responses in fear, we hypothesized that the DP may be playing a role in fear memory processes rather than in extinction. Indeed, in humans, the ventromedial prefrontal cortex has been implicated both in the inhibition and expression of fear ^14–17^. Nevertheless, despite this, the role for the DP in fear memory processing is largely unknown.

Here, we use a combination of whole-cell brain slice electrophysiology, activity-dependent neural tagging, immunohistochemistry, fiber photometry calcium imaging, and *in vivo* optogenetics to probe the role for the DP in the regulation of fear memory. We surprisingly found that rather than suppressing fear, neural ensembles in DP paradoxically function to encode fear memory and drive defensive behaviors. Our results provide a new role for the DP in fear memory encoding and fundamentally upend our understanding of the functional organization of the rodent mPFC.

## Results

### DP is activated in response to fear- but not extinction-related stimuli

To test whether the DP is involved in fear or extinction memory processes, we performed cFos immunohistochemical staining following exposure to three different behavioral paradigms. The first group of mice were subjected to tones only followed by a tone re-exposure test 24 hours later (**Figure 1A; Supplementary Figure 1**). The second group consisted of mice that were subjected to fear conditioning, which consisted of paired presentations of conditioned stimuli (CS; auditory tone) and unconditioned stimuli (US; foot shock) followed by a CS-evoked fear memory retrieval test 24 hours later (**Figure 1B; Supplementary Figure 1**). The third group consisted of mice subjected to paired fear conditioning followed by two consecutive days of CS extinction training. Animals from this group were then subjected to an extinction memory retrieval test 24 hours after the second day of extinction training (**Figure 1C; Supplementary Figure 1**). At 90 minutes after tone re-exposure or CS-evoked fear or extinction memory retrieval, mice were subjected to transcardial perfusion and brains were processed for cFos immunohistochemistry. cFos+ puncta were then quantified across the Cg1, PL, IL, and DP subregions of mPFC in mice from each behavioral group (**Figure 1D**). No differences in the density of cFos+ cells were observed across behavior groups in either the Cg1 or PL, consistent with our previous observations in the PL ^17^ (**Figure 1E**). In the IL, we observed a significant increase in the density of cFos+ cells in the extinction group compared to tones-only or fear conditioned mice (**Figure 1E**). This result is consistent with the role for the IL in extinction memory processing^18^. Compared to tones-only mice, those that were subjected to conditioning and fear memory retrieval exhibited a significantly higher density of cFos+ cells in the DP (**Figure 1E**). In contrast, mice subjected to extinction training exhibited a significantly lower density of cFos+ cells in the DP compared to conditioned mice.

**Figure 1.**
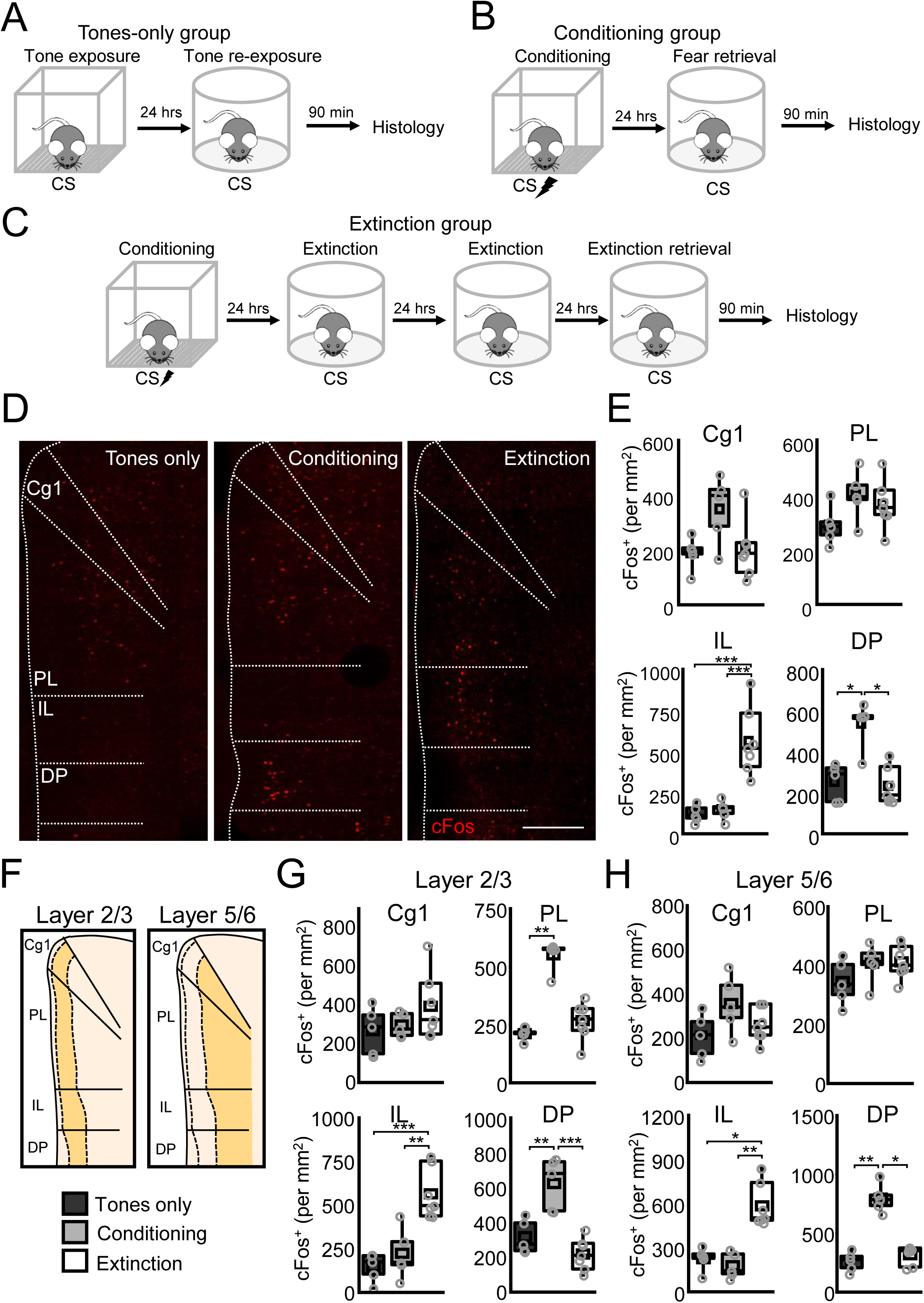
Fear conditioning drives activation of the DP. Mice were subjected either to (A) 6 tones (2 kHz, 20 s, 80 dB) followed by a tone re-exposure test consisting of 4 tone (2 kHz, 20 s, 80 dB) presentations 24 hours later; (B) paired presentations of a CS (2 kHz, 20 s, 80 dB) and US (2s, 0.7 mA) followed by a fear memory retrieval test consisting of 4 CS presentations (2 kHz, 20 s, 80 dB) 24 hours later; or to (C) paired presentations of a CS (2 kHz, 20 s, 80 dB) and US (2s, 0.7 mA) and 2 consecutive days of extinction training consisting of 20 CS (2 kHz, 20 s, 80 dB) presentations, and an extinction memory retrieval test consisting of 4 CS (2 kHz, 20 s, 80 dB) presentations. 90 minutes after tone re-exposure or fear or extinction memory retrieval, tissue was collected for cFos immunohistochemical analysis. (D) Representative histological images of prefrontal brain sections obtained from mice subjected to tones only, fear conditioning, or fear extinction. Scale = 500 µm. (E) Comparisons of cFos+ puncta density between tones only (n = 5 mice, 10 slices), conditioning (n = 5 mice, 10 slices), and extinction (n = 7 mice, 14 slices) groups in the anterior cingulate cortex 1 (Cg1: F_2,14_ = 4.08, P = 0.04, one-way ANOVA), prelimbic cortex (PL: F_2,14_ = 2.23, P = 0.14, one-way ANOVA), infralimbic cortex (IL: F_2,14_ = 19.34, P = 9.4 x 10^-4^, one-way ANOVA), and dorsal peduncular cortex (DP: Χ^2^ = 9.39 (2), P = 0.009, Kruskal-Wallis ANOVA). (F) Schematic cartoon of laminar demarcation across the mPFC. (G) Comparisons of cFos+ puncta density between tones only (n = 5 mice, 10 slices), conditioning (n = 5 mice, 10 slices), and extinction (n = 7 mice, 14 slices) groups in layer 2/3 of the anterior cingulate cortex 1 (Cg1: F_2,14_ = 1.49, P = 0.26, one-way ANOVA), prelimbic cortex (PL: X^2^ = 11.5 (2), P = 0.003, Kruskal-Wallis ANOVA), infralimbic cortex (IL: F_2,14_ = 17.9, P = 1.4 x 10^-4^, one-way ANOVA), and dorsal peduncular cortex (DP: F_2,14_ = 19.6, P = 8.8 x 10^-5^, one-way ANOVA). (H) Comparisons of cFos+ puncta density between tones only (n = 5 mice, 10 slices), conditioning (n = 5 mice, 10 slices), and extinction (n = 7 mice, 14 slices) groups in layer 5/6 of the anterior cingulate cortex 1 (Cg1: F_2,14_ = 2.69, P = 0.1, one-way ANOVA), prelimbic cortex (PL: Cg1: F_2,14_ = 1.81, P = 0.2, one-way ANOVA), infralimbic cortex (IL: Χ^2^ = 11.8 (2), P = 0.003, Kruskal-Wallis ANOVA), and dorsal peduncular cortex (DP: Χ^2^ = 10.8 (2), P = 0.005, Kruskal-Wallis ANOVA). *P < 0.05, **P < 0.01, ***P < 0.001, Tukey’s or Dunn’s post-hoc test. Box plots represent the median (center line), mean (square), quartiles, and 10-90% range (whiskers). Open circles represent data points for individual mice.

We next subdivided cFos counts across mPFC layers in each subregion (**Figure 1F**). In layer 2/3, we observed that there was a significant increase in the density of cFos+ cells in PL and DP following fear conditioning, as well as an increase in cFos in the IL following extinction training (**Figure 1G**). In layer 5/6, while we observed a similar increase in cFos in the IL and DP compared to layer 2/3, we did no observe either a fear- or extinction training-dependent increase in cFos in the PL (**Figure 1H**). These results suggest that rather than participating in extinction memory processing as was hypothesized previously, the DP may paradoxically be involved in fear-related memory processing.

### DP principal neurons exhibit fear learning-related plasticity

A hallmark of memory encoding is thought to be the expression of learning-related plasticity. We therefore sought to determine whether fear conditioning results in experience-dependent changes in DP neuron synaptic transmission. We performed whole-cell electrophysiological recordings in DP principal neurons (PNs) in acute brain slices prepared from wild-type mice. Prior to recording, mice were subjected to paired or unpaired fear conditioning (**Figure 2A; Supplementary Figure 2**). An additional group of mice were left trained and were only home cage-experienced. Acute brain slices were prepared 24 hours after behavioral training and spontaneous excitatory (EPSCs) and inhibitory (IPSCs) postsynaptic currents were recorded from DP PNs. Mice that were subjected to paired fear conditioning exhibited a higher frequency of spontaneous EPSCs onto DP PNs as compared to unpaired and naive control mice (**Figure 2B-C**). No group-dependent differences in spontaneous EPSC amplitude or IPSC frequency or amplitude were observed (**Figure 2D-E**).

**Figure 2.**
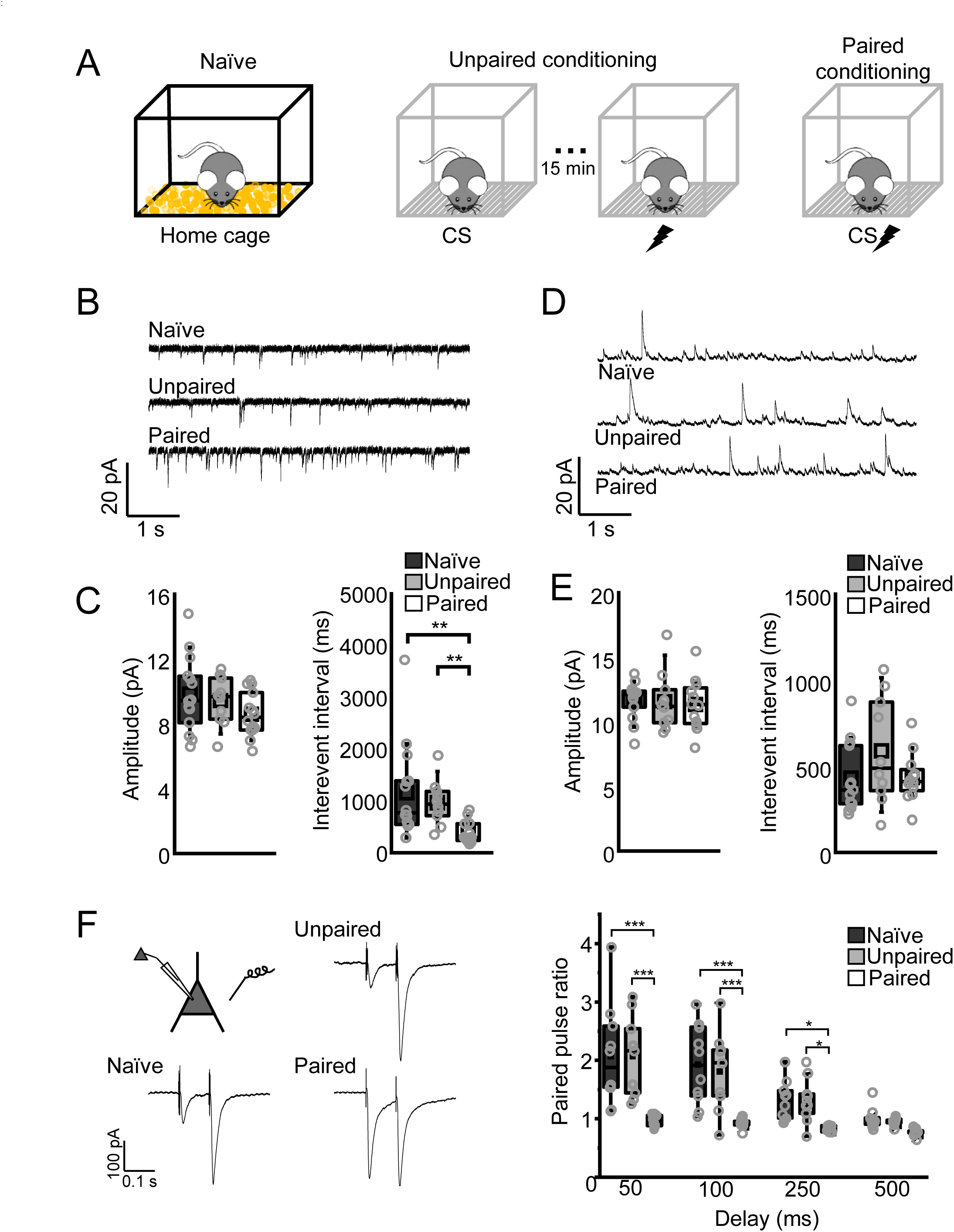
Fear conditioning drives increased glutamatergic transmission onto DP PNs. (A) Mice were either experienced to their home cage (naïve) or subjected to unpaired or paired presentations of a CS (2 kHz, 20 s, 80 dB) and US (2s, 1 mA). Whole-cell recordings were performed 24 hours after training. (B) Representative spontaneous excitatory postsynaptic current (EPSC) recordings in DP PNs from brain slices prepared from naïve (8 slices from 4 mice), unpaired (5 slices from 3 mice), and paired (8 slices from 4 mice) mice. (C) Quantification of the EPSC amplitude and interevent interval for naïve, unpaired, and paired groups. Amplitude: F_2,35_ = 1.67, P = 0.2, one-way ANOVA; naïve = 14 cells; unpaired = 10 cells; paired = 14 cells. Interevent interval: Χ^2^ = 14.32 (2), P = 7.77 x 10^-4^, Kruskal-Wallis ANOVA; naïve = 14 cells; unpaired = 10 cells; paired = 14 cells. (D) Representative spontaneous inhibitory postsynaptic current (IPSC) recordings in DP PNs from brain slices prepared from naïve, unpaired, and paired mice. (E) Quantification of the IPSC amplitude and interevent interval for naïve, unpaired, and paired groups. Amplitude: F_2,32_ = 0.084, P = 0.92, one-way ANOVA; naïve = 13 cells; unpaired = 10 cells; paired = 12 cells. Interevent interval: Χ^2^ = 2.2 (2), P = 0.33, Kruskal-Wallis ANOVA; naïve = 13 cells; unpaired = 10 cells; paired = 12 cells. (F) Cartoon schematic of recording configuration and representative excitatory paired-pulse recordings (100 ms delay) from naïve, unpaired conditioned, and paired conditioned mice. (G) Quantification of paired pulse ratio: F_6,48_ = 6.82, P = 2.83 x 10^-5^, interaction between training and delay, two-way repeated measures ANOVA, naïve = 10 cells; unpaired = 9 cells; paired = 10 cells. *P < 0.05, **P < 0.01, ***P < 0.001, Tukey’s or Dunn’s post hoc test. Box plots represent the median (center line), mean (square), quartiles, and 10-90% range (whiskers). Open circles represent data points for individual cells.

Similar patterns of plasticity were observed in recordings of miniature EPSCs and IPSCs (**Supplementary Figure 3**). Paired-pulse EPSC recordings from DP PNs revealed that paired fear conditioning results in a decreased paired-pulse ratio compared to naïve and unpaired groups (**Figure 2F**). Together, these results suggest that paired fear conditioning drives the potentiation of excitatory transmission onto DP PNs.

### DP but not IL neurons exhibit CS-related activity during conditioning and memory retrieval

The previous experiments lead to the observation that paired fear conditioning leads to strengthening of excitatory inputs onto DP PNs. These results may point towards a role for DP in cue-related fear memory processing. To test this possibility, we performed fiber photometry calcium imaging in the DP of mice undergoing fear conditioning and memory retrieval. Wild-type mice received unilateral infusions of GCaMP8f in DP as well as angled imaging fiber implantation immediately dorsolateral to the DP (**Figure 3A-B; Supplementary Figure 4**). We continuously recorded calcium associated transients from DP neurons during a pre-conditioning CS exposure test, during paired fear conditioning, and during CS-evoked fear memory retrieval. During fear conditioning, we observed a trial-dependent increase in cue-related activity, consistent with DP neurons having the ability to track and potentially encode information about the cue (**Figure 3C**). At 24 hours after conditioning, mice were placed in a neutral context and subjected to a cue-evoked fear memory retrieval test. Consistent with the idea that DP encodes cue-related information, we observed strong cue-elicited calcium signals. These signals were significantly larger than those recorded from the same mice in a preconditioning tone exposure test (**Figure 3D**). In contrast, we did not observe significant changes in calcium signals during spontaneous freezing bouts occurring during inter-CS intervals, suggesting that the DP does not signal freezing generally (**Supplementary Figure 5**). Moreover, DP recordings from mice that were subjected to unpaired conditioning followed by context retrieval revealed that DP activity is not related to contextual memory (**Supplementary Figure 6**). Together, these results suggest that the DP tracks and encodes cue-related information during fear conditioning and memory retrieval.

**Figure 3.**
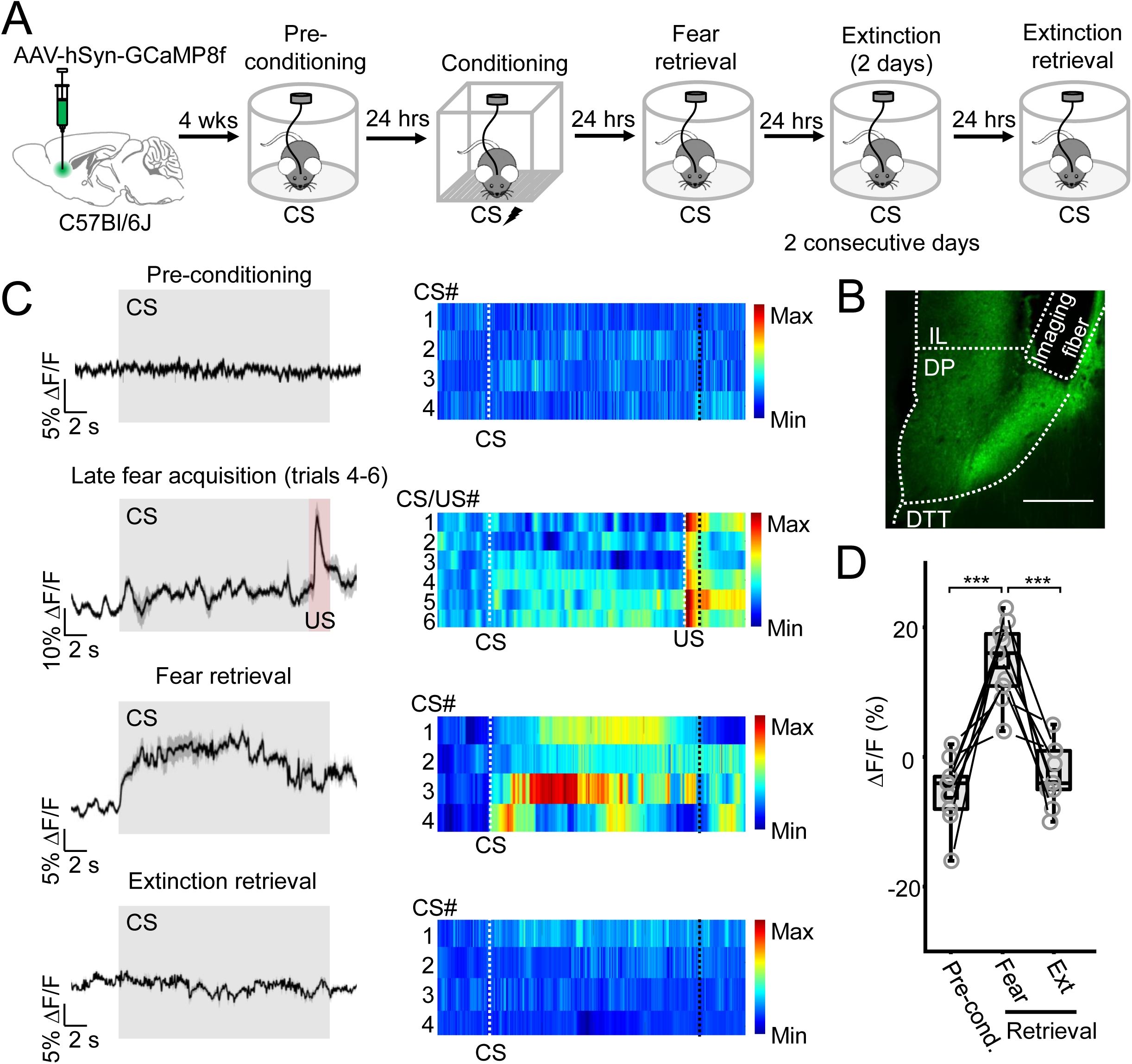
DP signals cue-related activity during fear memory acquisition and retrieval. (A) WT mice (n = 9) received infusions of a vector encoding GCaMP8f along with imaging fiber implantation directed at the DP. CS calcium-associated activity was monitored before, during, and after conditioning, in addition to after extinction. Fear conditioning consisted of a baseline period (300 s) followed by 6 pairings of CS (2 kHz, 20 s, 80 dB) and US (2 s, 0.7 mA). Pre-conditioning tone exposure, fear memory retrieval, and extinction memory retrieval all consisted of a baseline period (100 s) followed by four CS presentations (2 kHz, 20 s, 80 dB). (B) Representative histological image of GCaMP8f expression and fiber placement. Scale bar = 500 µm. (C) Left, raw calcium traces averaged across CS trials for each behavior test. Right, heat maps depicting individual calcium responses to each CS trial. (D) Comparison of ΔF/F percentage between pre- and post-conditioning CS presentation, as well as CS-associated activity following extinction training. F_2,15_ = 17.7, P = 1.13 x 10^-4^. **P < 0.01, Tukey’s post hoc test. Box plots represent the median (center line), mean (square), quartiles, and 10-90% range (whiskers). Open circles represent data points for individual mice. Lines represent data points from the same animal.

We also tested whether DP neurons may participate in extinction memory processing, as was previously proposed for this region ^8^. To test this possibility, 24 hours following the fear memory retrieval test, mice were subjected to two consecutive days of cue extinction training followed by an extinction memory retrieval test 24 hours later. In contrast to the signals observed during fear memory retrieval, we did not observe significant CS-related activity during cue presentation in the extinction retrieval test (**Figure 3C-D**). Instead, the calcium-associated activity returned to baseline-like levels that were not different from those recorded in the preconditioning tone-exposure test. Taken together, these results further support the idea that DP is involved in encoding cued fear, but not extinction memory processes.

The results thus far indicate that DP and IL likely play opposing roles in fear promotion and suppression respectively. We therefore further tested this idea *in vivo* by performing fiber photometry recordings in the IL. GcaMP8f was infused unilaterally and imaging fibers were implanted directed at IL. Mice were then subjected to the same behavioral testing as Figure 3 (**Supplementary Figure 7**). During fear conditioning and fear memory retrieval, we did not observe significant cue-related activity; these signals were not significantly different from those recorded during a pre-conditioning CS exposure test. In contrast, following extinction, we observed a significant increase in IL CS-related activity (**Supplementary Figure 7E and F**), consistent with our cFos experiments (**Figure 1**). These results indicate that DP and IL exhibit orthogonal activity patters in vivo and thus further suggest that they are functionally separate.

### DP neurons are activated by fear conditioning and reactivated in response to CS-evoked fear memory retrieval

The previous experiment revealed that DP neurons are activated both during fear memory acquisition and memory retrieval. However, it remained unclear whether the same or different DP neurons are activated during learning and retrieval, a pattern of activity that is thought to be a hallmark of memory encoding neurons. Towards this goal, we employed an activity-dependent neural tagging strategy that we previously used to tag fear-activated neurons ^19^. This strategy entails the use of a vector that allows for the activity-dependent expression of a 4-hydroxytamoxifen- (4-OHT) inducible Cre recombinase under the control of the synthetic enhanced synaptic activity-responsive element (ESARE-ERCreER) promotor ^20^. A vector encoding ESARE-ERCreER was infused in the the medial prefrontal cortex along with additional vectors encoding a Cre-dependent enhanced yellow fluorescent protein (DIO-eYFP) as well as an mCherry under the control of the human synapsin promotor (hSyn-mCherry) (**Figure 4A**). After 4 weeks, mice were subjected to either paired fear conditioning or to tones only (**Supplementary Figure 8**). Immediately after training, mice received intraperitoneal (IP) injections of either 4-OHT or vehicle. After two weeks, which is required for recombination and protein expression, mice were subjected to a CS-evoked fear memory retrieval test or tone re-exposure test in a neutral context after which they were transcardially perfused. Brain tissue was then processed for cFos immunohistochemistry (**Figure 4B**).

**Figure 4.**
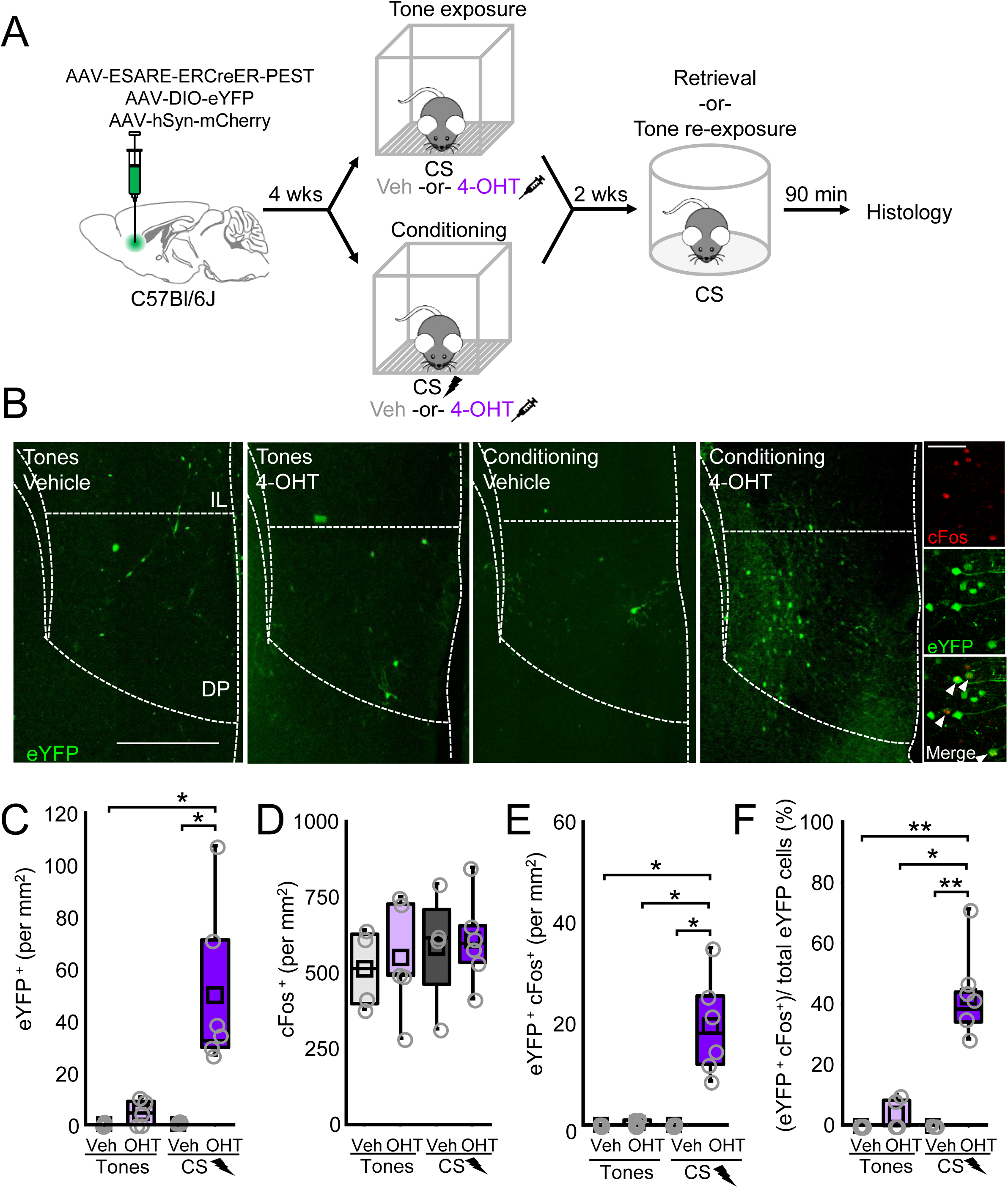
Learning-activated DP neurons are reactivated following cue-evoked memory retrieval. (A) WT mice received infusions of a cocktail of vectors encoding ESARE-ERCreER-PEST, a conditional eYFP, and mCherry into the DP. After 4 weeks, mice were subjected to tones only (2 kHz, 20 s, 80 dB) or to paired presentations of CS (2 kHz, 20 s, 80 dB) and US (2 s, 0.7 mA) and immediately injected with vehicle or 4-OHT after training. Following 2 weeks, mice were subjected to a CS-evoked fear memory retrieval test or tone re- exposure in a neutral context. Mice were subjected to transcardial perfusion 90 min later and tissue was used to stain against cFos. (B) Representative histological image from brain sections of tone-exposed and conditioned mice injected with vehicle or 4-OHT. Scale bar = 500 µm. Inset and arrowheads indicate tagged neurons staining positive for cFos (red). Scale bar = 75 µm. Comparisons between tones-only vehicle (n = 4 mice, 8 slices), tones-only 4-OHT (n = 5 mice, 10 slices), conditioned vehicle (n = 4 mice, 8 slices), and conditioned 4-OHT (n = 6 mice, 12 slices) for (C) eYFP+ (Χ^2^ = 13.75 (3), P = 0.003, Kruskal-Wallis ANOVA), (D) cFos+ (F_3,15_ = 0.288, P = 0.83, one-way ANOVA), (E) eYFP+ and cFos+ (Χ^2^ = 12.17 (3), P = 0.006, Kruskal-Wallis ANOVA), and (F) eYFP+ and cFos+/ eYFP+ (Χ^2^ = 15.31 (3), P = 0.0016, Kruskal-Wallis ANOVA). *P < 0.05, **P < 0.01, Tukey’s post hoc test. Box plots represent the median (center line), mean (square), quartiles, and 10-90% range (whiskers). Open circles represent data points for individual mice.

We observed that there were significantly more eYFP+ neurons in the DP of 4-OHT-injected conditioned mice compared to vehicle-injected conditioned mice or 4-OHT- or vehicle-injected tone-exposed mice (**Figure 4C**). While cFos-levels were comparable across groups (**Figure 4D**), there was an enrichment in the number of tagged neurons that were positive for cFos in the 4-OHT-injected conditioned group compared to those injected with vehicle or to vehicle- and 4-OHT-injected tones-only mice (**Figure 4E**). Comparisons of the proportion of cFos-positive tagged neurons revealed that 4-OHT-injected conditioned mice had significantly higher levels (∼40%) of fear retrieval-reactivated learning-tagged DP neurons compared to any other group (**Figure 4F**). These results reveal that ensembles of DP neurons exhibit activity patterns consistent with a role in fear memory encoding.

The results from our cFos (**Figure 1**) and fiber photometry experiments (**Figure 3**) suggested that DP neurons are activated in response to fear conditioning and cue-evoked fear memory retrieval, but not following extinction. To further test whether DP may be playing a role in extinction, we performed activity-dependent neural tagging following extinction training (**Supplementary Figure 9**). Following vector cocktail infusion in the IL and DP, mice were subjected to paired fear conditioning and three consecutive days of cue extinction training. Following the third extinction trial, mice received IP injections of either vehicle or 4-OHT. Two weeks later, brains were processed for immunohistochemistry. Consistent with our previous results, we only observed a small fraction of the total number of tagged neurons in the DP, and there was no difference in the number of tagged neurons in DP in the vehicle compared to the 4-OHT group. Importantly, a vast majority of tagged neurons instead resided in the IL, which has been implicated in extinction. Consistent with its role in extinction, there were significantly more tagged neurons in the IL of 4-OHT injected mice compared to those injected with vehicle. These results are also consistent with cFos immunohistochemistry results suggesting that cells in the IL but not DP are activated following extinction training (**Figure 1G-H**). Overall, these results provide additional evidence suggesting that the DP is involved in fear- but not extinction-related processes.

We additionally performed neural tagging in the DP following unpaired conditioning, in which mice were subjected to both the CS and US but in an explicitly unpaired manner (**Supplementary Figure 10**). Immediately after US delivery, mice received IP injections of either vehicle or 4-OHT. Two weeks after training, mice were subjected to a CS-evoked fear memory retrieval test and cFos immunohistochemistry was performed. Comparisons of the proportion of cFos+ eYFP neurons in vehicle versus 4-OHT-injected mice revealed no significant group differences. These results further support the idea that cue fear learning-tagged DP neurons represent a distinct learning-related population, rather than an ensemble activated by stress, contextual elements, or some other stimulus.

### Activity of fear-tagged DP neurons bidirectionally control fear memory expression

Given that learning-activated DP neurons are reactivated following CS-evoked fear memory retrieval, we next sought to test whether the activity of fear-tagged DP neurons can modulate fear memory expression. Towards this goal, we infused vectors encoding ESARE-ERCreER, Cre-dependent eYFP-tagged channelrhodopsin (DIO-ChR2-eYFP) or green fluorescent protein-tagged archaerhodopsin (FLEX-Arch-GFP), and hSyn-mCherry bilaterally into the DP of wild-type mice (**Figure 5A**). Following vector infusion, optic fibers were implanted bilaterally and directed at the DP (**Supplementary Figure 11**). After 4 weeks, mice were subjected to paired fear conditioning and immediately injected with vehicle or 4-OHT. After 2 weeks, mice were subjected to a fear memory retrieval test in a neutral context and laser stimulation-dependent alterations in freezing were assessed. In 4-OHT-injected mice expressing ChR2, optogenetic activation (473 nm, 20 Hz, 5 ms pulses, 20 s epochs) of fear-related DP neurons led to a significant increase in freezing over baseline (**Figure 5B**). Conversely, optogenetic silencing (532 nm, solid light, 20 s epochs) of fear-tagged DP neurons resulted in a reduction in CS-evoked freezing compared to CS presentation in the absence of optogenetic silencing (**Figure 5C**). Open field test during which optogenetic activation and silencing was performed revealed that there was no significant light-dependent effect on either locomotion or on anxiety-like behaviors (**Supplementary Figure 12**). Importantly, recordings from brain slices containing fear-tagged DP neurons expressing Arch-GFP or ChR2-eYFP revealed that the *in vivo* optogenetic manipulation procedures used result in reliable neural silencing or activation, respectively (**Supplementary Figure 13**). Overall, these results suggest that the activity of fear-tagged DP neurons bidirectionally controls fear memory expression.

**Figure 5.**
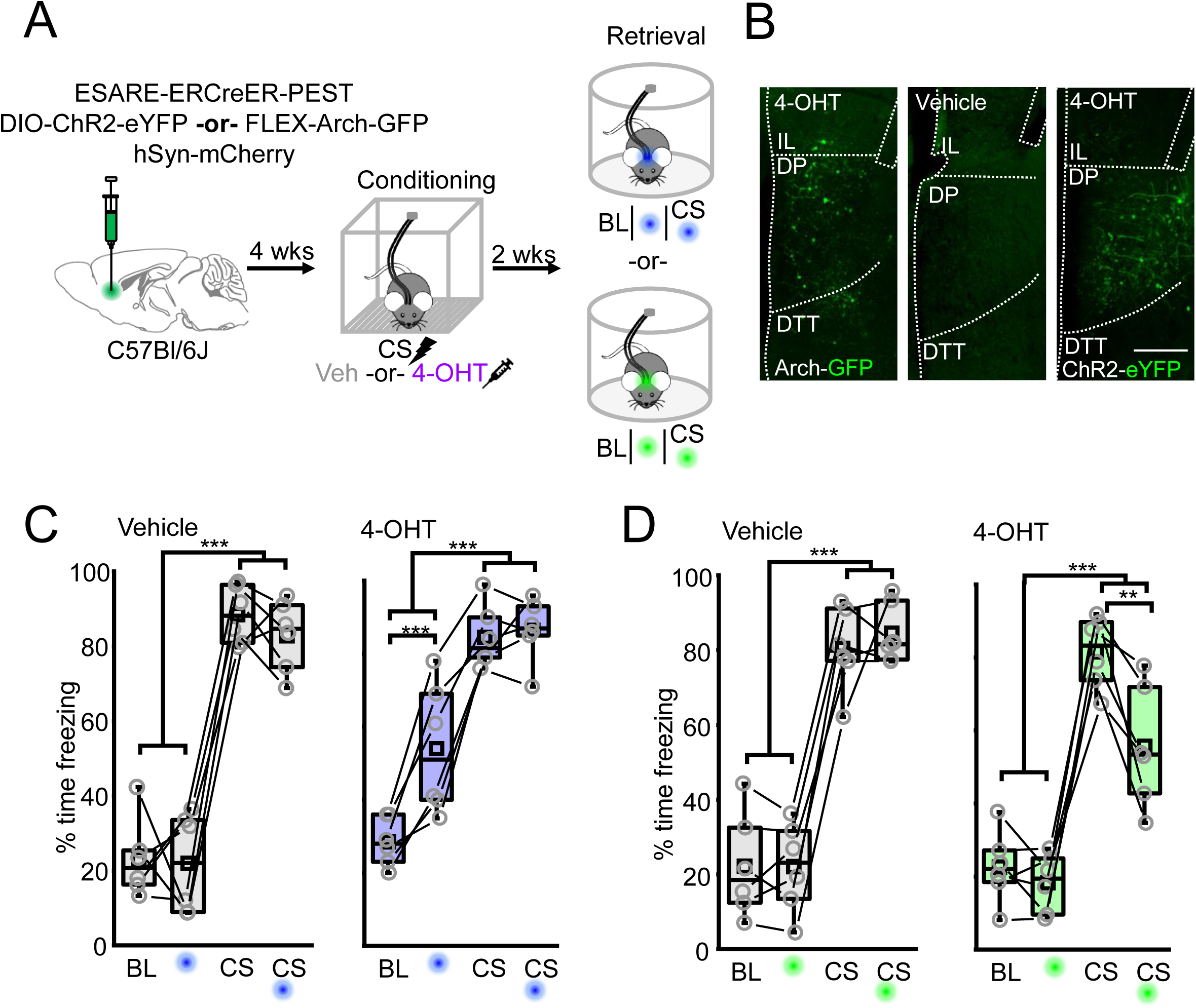
DP neuron activity bidirectionally impacts fear memory expression. (A) WT mice received infusions of vectors encoding ESARE-ERCreER-PEST, conditional ChR2-eYFP or conditional Arch-GFP, and mCherry along with implantation of optical fibers directed at the DP. Mice were subjected to paired fear conditioning and immediately injected with either vehicle or 4-OHT. After 2 weeks, mice were subjected to a CS-evoked fear memory retrieval in a neutral context with and without optogenetic manipulation. (B) Representative histological images from mice expressing Arch-GFP or ChR2-eYFP and receiving injections of vehicle or 4-OHT following fear conditioning. Scale = 400 µm. (C) Freezing quantified during CS (2 kHz, 20 s, 80 dB) and light (473 nm, 20 Hz, 10 ms pulses, 20 s epochs) presentation for animals expressing ChR2 in fear-tagged DP neurons. Vehicle: F_3,15_ = 22.14, P = 9.18 x 10^-6^, one-way repeated-measures ANOVA; n = 6 mice. 4-OHT: F_3,15_ = 73.39, P = 3.42 x 10^-9^, one-way repeated-measures ANOVA; n = 6 mice. (D) Freezing quantified during CS (2 kHz, 20 s, 80 dB) and light (532 nm, solid light, 20 s epochs) presentation for animals expressing Arch in fear-tagged DP neurons. Vehicle: F_3,15_ = 97.14, P = 4.73 x 10^-10^, one-way repeated measures ANOVA; n = 6 mice. 4-OHT: F_3,15_ = 53.29, P = 3.12 x 10^-8^, one-way repeated measures ANOVA; n = 6 mice. **P < 0.01; ***P < 0.001 Tukey post hoc test. Box plots represent the median (center line), mean (square), quartiles, and 10-90% range (whiskers). Open circles represent data points for individual mice. Lines represent data points from the same animal.

## Discussion

Here, by using multi-modal analyses, we reveal that the DP is involved in fear memory encoding and expression. We found that auditory fear conditioning but not extinction training drives activity of DP neurons. Moreover, fear conditioning potentiates glutamatergic transmission onto DP PNs compared to naive and unpaired littermate controls. By using fiber photometry recordings, we observed that DP neurons acquire CS-related activity during memory acquisition and that this activity persists during subsequent fear memory retrieval, but is lost after extinction training. Activity-dependent neural tagging following memory acquisition combined with cFos immunohistochemistry after retrieval indicated that ensembles of DP neurons are activated by learning and reactivated following memory retrieval. Finally, optogenetic activation of learning-tagged DP neurons induces fear memory expression whereas silencing significantly reduces cue-evoked freezing. Taken together, our results point towards an unexpected role for the DP in fear memory processing - an observation that is at odds with its proposed role in extinction ^8^.

The rodent vmPFC is most well-known for its role in memory extinction. In particular, lesion experiments ^21–23^, *in vivo* electrophysiological recordings and electrical stimulation ^24,25^, pharmacological microinfusion experiments ^26^, and *in vivo* optogenetic manipulations ^5^ have pointed towards a role for vmPFC in extinction memory processing, with a vast majority of this role attributed to the IL. However, some additional studies have provided conflicting results regarding this role. For example, *in vivo* pharmacological inhibition of vmPFC has paradoxically been shown to reduce the expression of conditioned fear ^27^. Moreover, additional lesion studies found that vmPFC lesions had no effect on extinction memory acquisition or retrieval ^28,29^. These conflicting findings are also reflected in the human literature where studies have implicated the vmPFC in extinction ^1,14,15^ while other studies have suggested vmPFC activity contributes to the pathology of post-traumatic stress disorder ^16,17^. While the conflicting results in rodents can potentially be explained by compensatory mechanisms or differences in experimental execution, one additional possibility may be due to differences in the lesion/ infusion locus within vmPFC. The results of our current study that implicate the DP in fear but not extinction memory processes may potentially account for the conflicting findings in the field.

While our findings have provided numerous lines of evidence suggesting the DP plays a role in fear memory encoding and retrieval but not extinction, a recent study by Botterill and colleagues reported that the DP/DTT seems play a role in fear extinction rather than fear encoding and retrieval^9^. These differences may relate to which specific DP outputs were manipulated, the method of manipulation, and anatomical differences in viral targeting. Moreover, an additional study revealed sexual dimorphic activity profiles in the DP following auditory fear conditioning, where female mice exhibited significantly higher levels of conditioning-induced cFos expression compared to males^30^. While most of our experiments are not sufficiently powered to perform rigorous sex differences analyses, our results do not point toward any gross differences in DP activity in male and female mice. Whether conditioning engages distinct cell types or circuits within the DP of male and female mice remains to be an important area for future investigation.

During the revision of this manuscript, an additional study was published that points towards a role for DP excitatory projections to the central amygdala in the control of fight or flight^13^. The authors observed that neurons in layer 2/3 project to central amygdala whereas those in layer 5/6 project to the dorsal medial hypothalamus, a circuit implicated in mediating sympathetic responses to psychosocial stress^10^. As we observed increased cFos expression across DP cortical layers (**Figure 1**), the population we examined is likely comprised of neurons projecting to both of these downstream targets. Indeed, sympathetic responses and circuits of fight and flight are both central in fear memory encoding processes. Future studies examining the specific projection profiles of fear-tagged DP neurons will be needed to determine other targets of this population.

As the role for DP in fear memory encoding and retrieval as well as in processing other emotional stimuli continues to be investigated, additional questions remain to be answered in future studies. One such question is why might both the PL and the DP function to encode fear memory? Are each of these brain regions encoding distinct aspects of the fear memory? Similar to the DP, PL is activated both during fear memory encoding and retrieval^31,32^, and its activity drives the expression of defensive behaviors^25,26,33,34^. Despite their similar function in promoting fear, there are also some notable differences in fear memory processes in DP and PL. In contrast to the PL, which is involved in both cued and contextual fear^21,33,35^, DP neurons signal cued and not contextual fear memory (**Supplementary Figure 6** and 10). Moreover, PL neurons exhibit tone responses in naïve mice^32,36^, whereas the same responses do not exist in the DP, at least at the level of our analyses. Insights to the circuits that underlie these differences may be gained through the systematic investigation of projection pathways that differentially engage the PL versus the DP. Indeed, functional *in vivo* oscillation measurements across mPFC subregions in anesthetized rats have revealed vast differences in the PL and DP, suggesting that these regions likely have unique functions and/or connectivity patterns^37^. Moreover, an anatomical analysis reveals bidirectional connectivity between the PL and DP^38^, suggesting that they have the capacity to convey information to each other during memory encoding. Therefore, it will be important for future studies to reveal the fundamental connectivity of long-range inputs to DP to gain insight to these questions.

Our results suggest that a relatively sparse ensemble of DP neurons functions to encode fear memory. While we have characterized the functional contributions of this ensemble, several questions remain. First, it remains unknown what neuron type(s) comprise this ensemble and how these distinct cell types contribute to memory encoding. Our previous work revealed that somatostatin-expressing interneurons (SST-INs) comprise a relatively large part of the fear-activated ensemble in the PL^19^. Whether these or another subset of interneurons are recruited into the DP ensemble remains unknown. Additionally, future studies should focus on revealing the local microcircuit connectivity and plasticity mechanisms that support the recruitment of the DP ensemble. Indeed, our previous work in the PL revealed that there is extensive rebalancing of microcircuit activity that resulted in SST-IN-mediated inhibition of parvalbumin-expressing interneuron (PV-IN) activity, driving the disinhibition of PNs ^39^. Whether this circuit organization and plasticity mechanism is conserved in the DP is unknown and should be studied in future work.

Overall, our results provide critical insights to fear memory encoding and expression in the mouse DP. These findings open several new areas of investigation and fundamentally redefine our understanding of the functional landscape within the rodent mPFC.

### Limitations of the study

Although our study provides several lines of evidence suggesting that the DP encodes fear memory, there are some limitations that are important to consider. First, although we performed electrophysiological analyses that revealed that DP PNs undergo fear-related plasticity, we did not identify and test the causal contribution of potential sources of the plasticity. This will be important for identifying how the DP is engaged during fear conditioning, and how it is incorporated into our existing understanding of fear circuitry. A second limitation lies in our fiber photometry experiments. Since fiber photometry only reports on bulk calcium-associated neural activity, it is possible that a small population of extinction-related DP neurons exists, but that their activity is masked by a larger or more active population of fear-encoding neurons. Since this population, if it exists, would likely be small, it will be important to utilize experimental approaches that enable single-cell resolution such as silicon probe or microendoscopic recordings, to examine contributions of this potential population. Importantly, however, these limitations do not impact our interpretation that the DP encodes fear memory.

## Author contributions

K.A.C. and R.L.C initiated the project. K.A.C supervised the research. K.A.C, R.C.C, Z.R.D, B.L.F., and R.L.C. designed experiments. K.A.C., R.C.C., Z.R.D., B.L.F., A.M., J.D., and V.A.L. performed the research and data analysis. K.A.C., R.C.C., Z.R.D., and B.L.F. wrote the manuscript.

## Acknowledgments

This work was supported by the National Institute of Mental Health grant no. R00 MH122228 to K.A.C., Brain and Behavior Research Foundation Young Investigator Award to K.A.C., Joe W. and Dorothy Dorsett Brown Foundation grant to K.A.C., James A Pittman, Jr., MD Scholar Award to K.A.C., and T32 GM135028 to B.L.F. We thank Dr. Sofia Beas for valuable comments on the manuscript. We also thank members of the Cummings lab for feedback on the manuscript and experiments. Graphical abstract was made with BioRender.

## STAR★Methods

### Resource availability

#### Lead contact

Further information and requests for resources and reagents should be directed to and will be fulfilled by the lead contact, Kirstie Cummings.

#### Materials availability

This study did not generate new unique reagents.

#### Data and code availability

- All data reported in this paper will be shared by the lead contact upon request.
- This paper does not report original code.
- Any additional information required to reanalyze the data reported in this work paper is available from the lead contact upon request.

### Experimental model and study participant details

#### Animals

The use of animals in all experimental procedures was approved by the Institutional Animal Care and Use Committee at the Heersink School of Medicine at the University of Alabama at Birmingham. Both male and female mice at the age of postnatal day 42 (P42) were used for stereotaxic surgery and behavioral experiments were then performed at P60-90. Mice used in experiments were the C57Bl/6J genotype (stock no. 000664). Animals were housed in groups of 2-6 with a 12 h light-dark cycle and *ad libitum* access to food and water. Equal numbers of males and females were used for all experimental groups.

### Method details

#### Behavioral training paradigms

Paired auditory fear conditioning was performed by subjecting mice to 6 pairings of a pure auditory tone (CS; 80 dB, 2 kHz, 20 s) with a co-terminating scrambled foot shock (US; 0.7- 1.0 mA, 2 s). Unpaired conditioned mice were subjected to 6 CS presentations after which they were returned to their home cage for 15 min and then placed back into the arena during which they received 6 US presentations. Naive mice only experienced the home cage. Tones-exposed mice were placed in the conditioning arena and subjected to 6 CS tones in the absence of the US. For extinction training, 24 hours after paired fear conditioning, mice were placed in a neutral context (context B) consisting of distinct visual, olfactory, and tactile cues and subjected to 20 CS tones. An additional extinction session was performed the next day for a total of 2 consecutive days. For either fear or extinction memory retrieval in cFos, electrophysiological, and calcium imaging experiments (Figures 1, 2, and 3), mice were placed in context B and subjected to presentation of 4 CS tones at 24 hours after either fear conditioning or the second day of extinction training, respectively. Tone re-exposure was performed in context B by presenting animals 4 CS tones at 24 hours after tone exposure. For fear memory retrieval or tones re-exposure following neural tagging (Figures 4 and 5), mice were placed in context B and subjected to presentation of 4 CS tones at 2 weeks (due to limitations of vector expression timeline) after fear conditioning or tone exposure, respectively. Littermates were used for each experiment whenever possible and were randomly distributed across behavior groups in an interleaved fashion.

#### Immunohistochemistry

Immunohistochemistry was performed for cFos and/or GFP. 90 minutes after the last behavioral test, after deep isoflurane anesthesia was induced, mice were transcardially perfused with phosphate buffered saline (PBS; pH 7.4) followed by 4% paraformaldehyde in PBS (PFA; pH 7.4) before removal of the brains. Brains were then fixed in 4% PFA overnight before sectioning 50 µm coronal slices using a Leica VT1000S vibratome (Leica Biosystems, Deer Park, IL, USA).

Brain sections were blocked with 5% normal goat serum in PBS + 0.3% Tween-20 for one hour at room temperature. Afterwards, slices were incubated at 4° C overnight with rabbit anti-cFos primary antibody (1:1000; Millipore ABE457). The following day, slices were incubated for 2 hours with a secondary goat anti-rabbit antibody conjugated to Alexa 647 (1:500; Jackson Immunoresearch, 111-605-045) with 5% normal goat serum in PBS + 0.3% Tween-20.

For experiments requiring staining against GFP, slices were then incubated overnight with a chicken anti-GFP primary antibody (1:500; Millipore AB16901) with 5% normal donkey serum in PBS + 0.3% Tween-20. The next day, slices were incubated with a secondary donkey anti-chicken antibody conjugated to Alexa 488 (1:500; Jackson Immunoresearch, 703-095-155) for 2 hours at room temperature.

Following immunohistochemical staining, slices were mounted with ProLong Gold antifade reagent with DAPI (Life Technologies, Eugene, OR, USA). Imaging was performed using an upright Echo Revolution hybrid microscope (BICO, San Diego, CA, USA) or a Nikon Eclipse Ni-E microscope (Nikon Instruments, Melville, NY, USA). Cell counting for cFos or/ and GFP labeled cells was performed with the Cell Counter plug-in through the ImageJ program (NIH, Bethesda, MD, USA). Quantification was performed for either or both cFos+ and GFP+ cells, as well as for overlapping cFos+/GFP+ cells by experimenters blind to experimental conditions and groups.

Anatomical demarcations were determined by creating overlays aligned to the *Paxinos and Franklin’s The Mouse Brain In Stereotaxic Coordinates* atlas^40^ in Photoshop. These overlays were then imported into ImageJ (NIH, Bethesda, MD, USA) for analysis. Overlays were created by experimenters blind to experimental conditions, and cFos expression, tagged neuron distribution, etc. were not used as a factor in determining anatomical borders. In an effort to enhance robustness and reliability of data, several blinded experimenters performed these demarcations across multiple cohorts of mice.

#### Whole-cell brain slice electrophysiology

Mice were subjected to paired fear conditioning, unpaired fear conditioning, or were naive. 24 hours after training, mice were used for electrophysiology recordings.

Mice were deeply anesthetized by way of isoflurane administration prior to decapitation. Brain slices containing the mPFC were sectioned at 300 µm on a Leica VT1200S vibratome in chilled carbogen-bubbled low sodium sucrose cutting solution containing (in mM) 210 sucrose, 11 glucose, 26.2 NaHCO_3_, 2.5 KCl, 1 NaH_2_PO_4_, 4 MgCl_2_, 0.5 CaCl_2_, and 0.5 ascorbate. Slices were then recovered for 45 min in warmed (32° C) carbogen-bubbled normal artificial cerebrospinal fluid containing (in mM) 119 NaCl, 26.2 NaHCO_3_, 11 glucose, 1 NaH_2_PO_4_, 2 MgCl_2_, 2.5 KCl, 2 CaCl_2_. Slices were kept at room temperature. Whole-cell electrodes (3-4 MΩ) were fabricated from fire polished borosilicate capillaries and filled with internal solution containing (in mM) 120 Cs-methanesulfonate, 10 HEPES, 10 Na-phosphocreatine, 1 QX-314, 8 NaCl, 0.5 EGTA, 4 Mg-ATP, 0.4 Na-ATP (pH 7.25; 290-300 mOsmol). Cells were visualized on an upright microscope equipped with differential interference contrast optics. Principal neurons were identified visually (large pyramidal somas) and electrophysiologically (low membrane resistance (<75 MΩ) and high capacitance (>100 pF). Cells were sampled across superficial and deep layers and pooled.

Spontaneous excitatory (EPSC) and inhibitory (IPSC) postsynaptic currents were recorded in standard artificial cerebrospinal fluid (ACSF) by clamping cells at -60 mV or 0 mV, respectively. Miniature EPSCs and IPSCs were recorded in standard ACSF with the addition of 1 µM tetrodotoxin (TTX). Each EPSC and IPSC recording lasted at least 5 min. Paired pulse measurements were performed with a bipolar stimulating electrode placed adjacent to recorded cells. Averages of 7-10 sweeps per cell per delay were used for calculating paired pulse ratio. Paired pulse ratio was calculated by dividing the amplitude of the second current by the amplitude of the first. Data were sampled at 10 kHz and low-pass-filtered at 3 kHz using Multiclamp 700B amplifier (Molecular Devices, version 2.2.2.2). Data were analyzed in Easy Electrophysiology (version 2.6.1) by experimenters blind to experimental groups. Access resistance and leak current were monitored throughout each recording. Recordings that yielded unstable currents (>150 pA leak; >20% change in access resistance) or which did not meet cell type characterization requirements were excluded. Cell exclusion was rare and occurred <2% of the time.

#### Stereotaxic vector infusion and fiber implantation

AAV2/8-ESARE-ERCreER-PEST was received as a gift from Dr. Harihiko Bito (U. Tokyo) and custom packaged at Boston Children’s Hospital Vector Core. AAV1-DIO-eYFP (Cat. No. 27056), AAV8-hSyn-mCherry (Cat. No. 114472), AAV1-hSyn-GCaMP8f (Cat. No. 162376), AAV1-DIO-hChR2(H134R)-eYFP-WPRE (Cat. No. 20298), and AAV1-FLEX-Arch-GFP (Cat. No. 22222) were purchased from Addgene. For all experiments, vector infusions occurred at P42. For activity-dependent tagging quantification experiments, vectors were infused bilaterally in the DP (200 µl; A/P: 1.55; M/L: +/- 0.25; D/V: -3.7) or in DP and IL (400 µl; A/P: 1.55; M/L: +/- 0.25; D/V: -3.7-3.5). Vectors were mixed at a 2 (ESARE-ERCreER-PEST) to 7 (DIO-eYFP) to 1 (hSyn-mCherry) ratio just prior to stereotaxic infusion. For fiber photometry experiments and *in vivo* optogenetics, vectors were infused unilaterally (fiber photometry) or bilaterally (*in vivo* optogenetics) into the DP (A/P: 1.55; M/L: + and/or – 2.3; D/V: -4.1; 32°∡) or IL (A/P: 1.55; M/L: + and/or - 1.25; D/V: -2.7; 15°∡). For fiber photometry experiments, an imaging fiber (400 µm diameter, 0.48 NA, Doric Lenses) was implanted immediately above the DP (A/P: 1.6; M/L: +/- 1.65; D/V: -3.9; 32°∡) or IL (A/P: 1.55; M/L: + and/or - 1.25; D/V: -2.6; 15°∡). For *in vivo* optogenetic experiments, optic fibers were implanted directed at DP (A/P: +1.6; M/L: +/- 1.65; D/V: -3.8; 32° angle). Vector incubation times were previously determined ^19,39^ or were determined empirically for this study.

#### Activity-dependent neural tagging and cFos immunofluorescence

At 3.5 weeks post vector cocktail infusion, mice were handled for three days consecutively prior to behavior. Mice were subjected either to auditory paired fear conditioning, tones-only, or unpaired fear conditioning. For mice used to tag extinction-related mPFC neural ensembles, animals were subjected to extinction training for 3 consecutive days starting at 24 hours after fear conditioning. Neural tagging was performed through an intraperitoneal (IP) injection of 4-OHT (10 mg/kg) or vehicle solution immediately after the completion of the last day of training, as was done previously ^19^. Two weeks after training and activity-dependent tagging, mice were subjected to a fear memory retrieval test in a neutral context (context B). Mice were sacrificed 90 minutes post-retrieval for cFos and GFP immunohistochemistry as described above.

#### Fiber photometry recording and analysis

Fiber photometry recordings were performed by using a complete system from Tucker-Davis Technologies, as we have done previously ^39^. At 4 weeks after vector infusion and imaging fiber implantation, mice were habituated to handling and patch cord tethering for 3 consecutive days. Mice were placed in a neutral arena with white floor and rounded wall inserts (context B) and subjected to 4 CS presentations (CS: 80 dB, 2 kHz, 20 s). 24 h later, mice were subjected to either a paired or unpaired fear conditioning in an arena with a metal grid floor and square metal walls (context A). Mice were then subjected to a fear memory retrieval test at 24 h post-conditioning. 24 hours later, mice were subjected to two consecutive days of extinction training in context B. At 24 h post extinction training, mice were subjected to an extinction memory retrieval test in context B. Calcium signals were sampled at 6 kHz and continuously recorded throughout the duration of each behavioral test. Transistor-transistor logic (TTL) signals were generated through MedAssociates software and were used to produce timestamps for the start and stop of each session as well as for CS presentation epochs. Prior to the start of each session, calcium signals were recorded for a duration of 2 min to ensure stable recordings.

Data were processed and analyzed using custom-written code from the Tucker-Davis Technologies website as described previously ^39^. Briefly, CS-associated activity changes were calculated by normalizing the calcium signal during the 20 s CS presentation to a 20 s baseline period occurring immediately prior to CS onset. The percentage change in fluorescence (%ΔF/F) was calculated by subtracting the peak baseline signal from the peak signal during CS presentation and dividing by the peak baseline signal (%ΔF/F = (F_peak_CS - F_peak_BL)/F_peak_CS)). Reported values represent within-animal changes. At the conclusion of experiments, mice were subjected to transcardial perfusion with PBS followed by 4% paraformaldehyde and imaging fiber placement and GCaMP8f expression was assessed. Mice with incorrect fiber placements and/or GCaMP8f expression were excluded from analyses.

#### *In vivo* optogenetics

At 3.5 weeks after vector infusion and optic fiber implantation, mice were habituated to handling and tethering to patch cords for 3 consecutive days. Mice were subjected to paired fear conditioning and then received intraperitoneal injections of either 4-OHT (10 mg/kg) or vehicle immediately after training and returned to their home cage. After two weeks, mice were subjected to a CS-evoked fear memory retrieval test in a neutral context (context B). Optogenetic activation (473 nm, 20 Hz, 5 ms pulses, 20 s epochs) or silencing (532 nm, solid light, 20 s epochs) was performed in animals expressing ChR2 or Arch, respectively. Light intensities delivered to the brain were measured and calculated prior to each experimental day and ranged from 6-8 mW. Optogenetic manipulations were performed alone or in combination with the CS in an alternating manner (e.g. Baseline: light-on, light-off; CS: CS + light, CS alone, etc) and the order or light presentation was counterbalanced. Each epoch (baseline, light-on; baseline, light-off; CS, light-on; CS, light-off) was repeated twice per animal and the average was reported. Videos were scored by experimenters blind to experimental conditions and groups. At the end of the experiment, mice were subjected to transcardial perfusion and brains were sectioned for immunohistochemical analysis to confirm correct optic fiber placement and vector expression. Mice with incorrect fiber placement and/or viral expression were removed from analyses.

#### Open field test

Mice from the *in vivo* optogenetics experiments (Figure 5) were also subjected to optogenetic stimulation in an open field. The test was 10 minutes in duration, and consisted of alternating and counterbalanced epochs of 1 minute light-on and 1 minute light-off periods. The open field measured 40 cm length x 40 cm width x 40 cm height and tracked movement via infrared beam breaks (MedAssociates, Fairfax, VT). Average distance moved and average percent time spent in the margin space was calculated and reported for each mouse for the light-on/ light-off epochs.

#### Electrophysiological validation of *in vivo* optogenetic manipulations

To validate the modes of optogenetic manipulation of fear-related DP neurons used *in vivo* (Figure 5), mice that received infusions of vectors encoding ESARE-ERCreER and either DIO-ChR2-eYFP or FLEX-Arch-GFP were subjected to paired fear conditioning and intraperitoneal injection of 4-OHT (10 mg/kg). After 2 weeks, mice were used to prepare brain slices, as described above. DP neurons expressing ChR2 or Arch were targeted in slice via fluorescence. For DP neurons expressing Arch, after achieving whole-cell configuration, slow current was injected to induce spontaneous action potential firing. Green light (532 nm; solid light; 8 mW; 20 sec epochs) was delivered via LED-coupled objectives for a total of 4-5 times per cell. The average firing frequency and magnitude of hyperpolarization was calculated during light off and light on epochs. For DP neurons expressing ChR2, after achieving the whole-cell configuration, pulses of blue light (473 nm; 20 Hz, 5 ms pulses, 20 sec epochs) were delivered via LED-coupled objectives and action potential firing was recorded.

### Quantification and statistical analysis

Statistical testing and graph plotting were performed using OriginPro 2019 version 9.6.0.172. Levine’s and Shapiro-Wilk tests were performed prior to parametric statistical testing to test whether data exhibited homogeneity of variance and normal distribution, respectively. If either or both conditions were not met, data were subjected to non-parametric statistical analyses. All statistical details, including statistical tests used, the exact value of each n, and what n represents can be found in each Figure or Supplemental Figure legend. In all Figures and Supplementary Figures, box plots represent the median (center line), mean (square), quartiles, and 10-90% range (whiskers). Open circles represent individual data points, which is defined in each legend (e.g., mice, cells, etc.).

## Supplemental figure titles

Supplementary Figure 1, related to Figure 1. Behavior analysis for mice used for cFos immunohistochemistry.

Supplementary Figure 2, related to figure 2. Behavior analysis for mice used for electrophysiology recordings.

Supplementary Figure 3, related to figure 2. Fear conditioning drives increased glutamatergic transmission onto DP PNs.

Supplementary Figure 4, related to figure 3. Fiber photometry fiber placements and behavioral data.

Supplementary Figure 5, related to figure 3. Fiber photometry measurements during spontaneous inter-CS freezing bouts.

Supplementary Figure 6, related to figure 3. Fiber photometry measurements in DP during unpaired fear conditioning and contextual exposure.

Supplementary Figure 7, related to figure 3. IL signals cue-related activity during extinction memory retrieval.

Supplementary Figure 8, related to figure 4. Viral spread and behavior analysis associated with neural tagging data.

Supplementary Figure 9, related to figure 4. Behavior analysis and histology for neural tagging in response to extinction training.

Supplementary Figure 10, related to figure 4. Behavior analysis and histology for neural tagging in response to unpaired conditioning.

Supplementary Figure 11, related to figure 5. Fiber placements, viral expression, and behavioral data for optogenetic experiments.

Supplementary Figure 12, related to figure 5. Optogenetic manipulations during the open field test.

Supplementary Figure 13, related to figure 5. Electrophysiological validation of optogenetic manipulations.

---

**Supplementary Figure 1, related to Figure 1.**
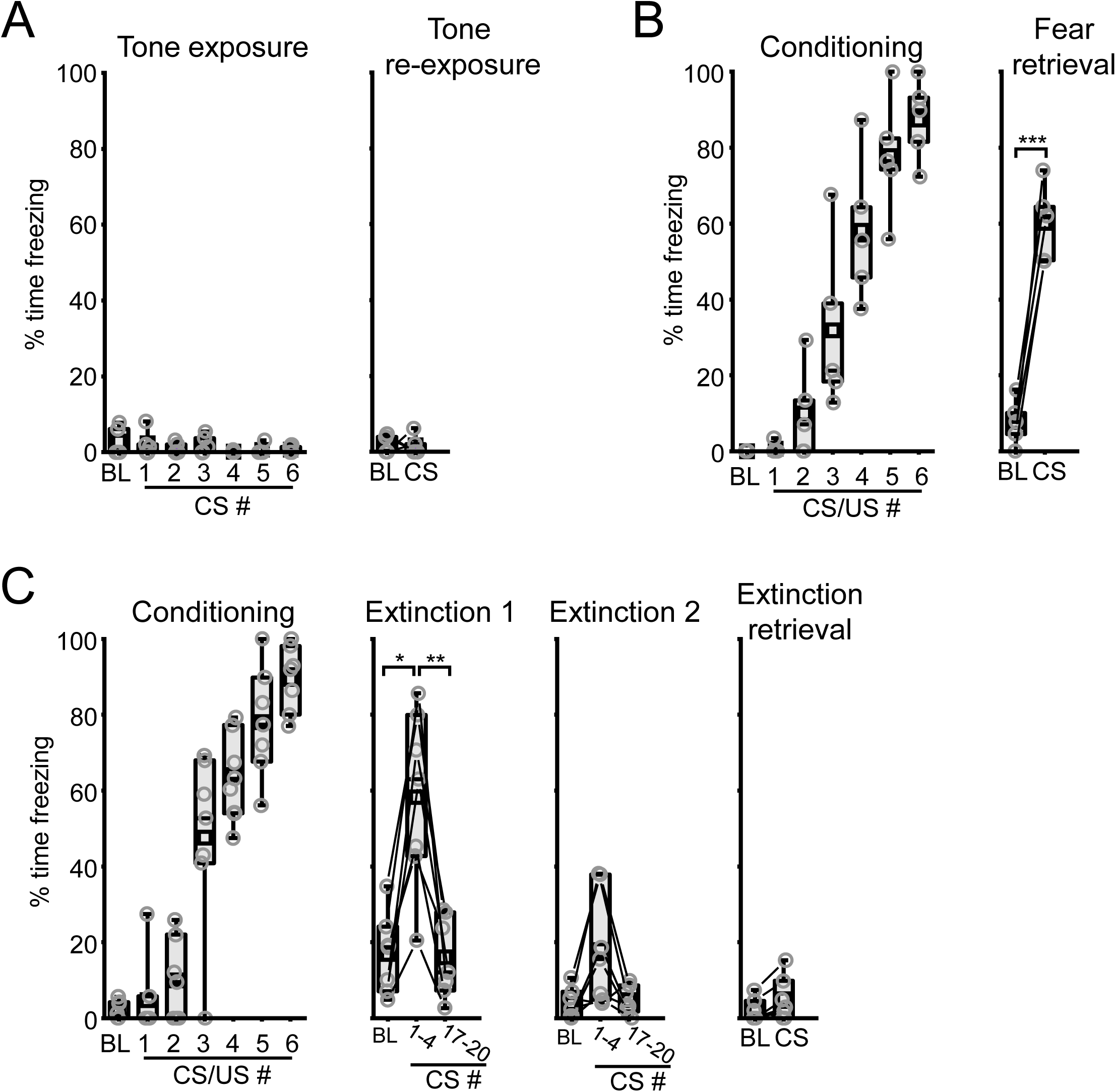
Behavior analysis for mice used for cFos immunohistochemistry. (A) Quantification of freezing for mice subjected to tone exposure and tone re-exposure (W = 5, P = 1, Wilcoxen signed rank test). (B) Quantification of freezing for mice that were fear conditioned and subjected to fear memory retrieval (t(4) = -11.52, P = 3.24 x 10-4, Paired t-test). (C) Quantification of freezing for mice that were fear conditioned and subjected to a first day of extinction training (X^2^ (2) = 10.57, P = 0.005, Friedman ANOVA), a second day of extinction training (X^2^(2) = 5.43, P = 0.066, Friedman ANOVA), and extinction memory retrieval (W = 2, P = 0.094, Wilcoxen signed rank test). *, P < 0.05, **, P < 0.01, Dunn’s post-hoc test. Box plots represent the median (center line), mean (square), quartiles, and 10-90% range (whiskers). Open circles represent data points for individual mice.

**Supplementary Figure 2, related to figure 2.**
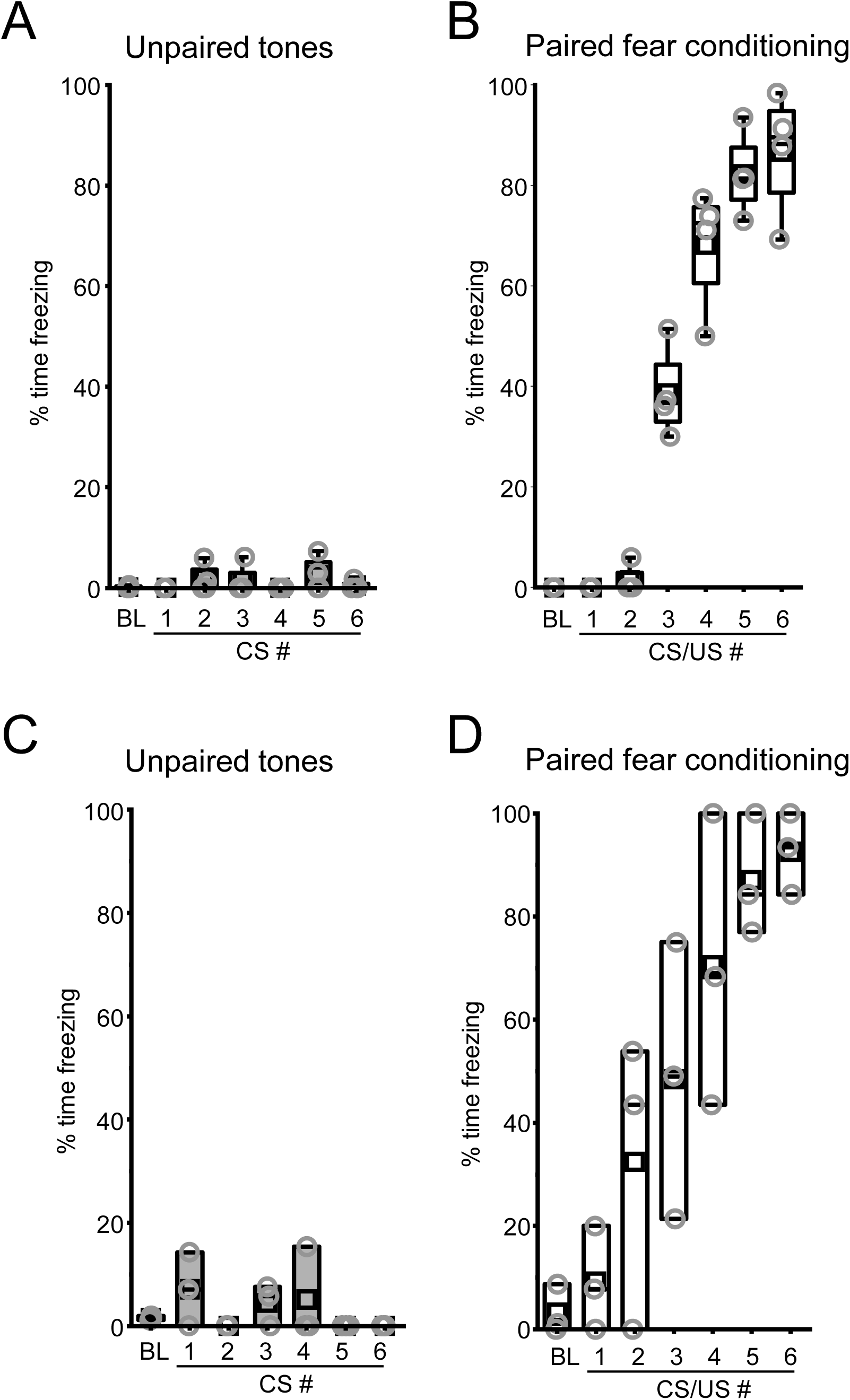
Behavior analysis for mice used for electrophysiology recordings. Quantification of freezing during unpaired tone presentation for mice used for (A) spontaneous synaptic recordings and (C) paired pulse recordings. Quantification of freezing during paired fear conditioning for mice used for (B) spontaneous synaptic recordings and (D) paired pulse recordings. Box plots represent the median (center line), mean (square), quartiles, and 10-90% range (whiskers). Open circles represent data points for individual mice.

**Supplementary Figure 3.**
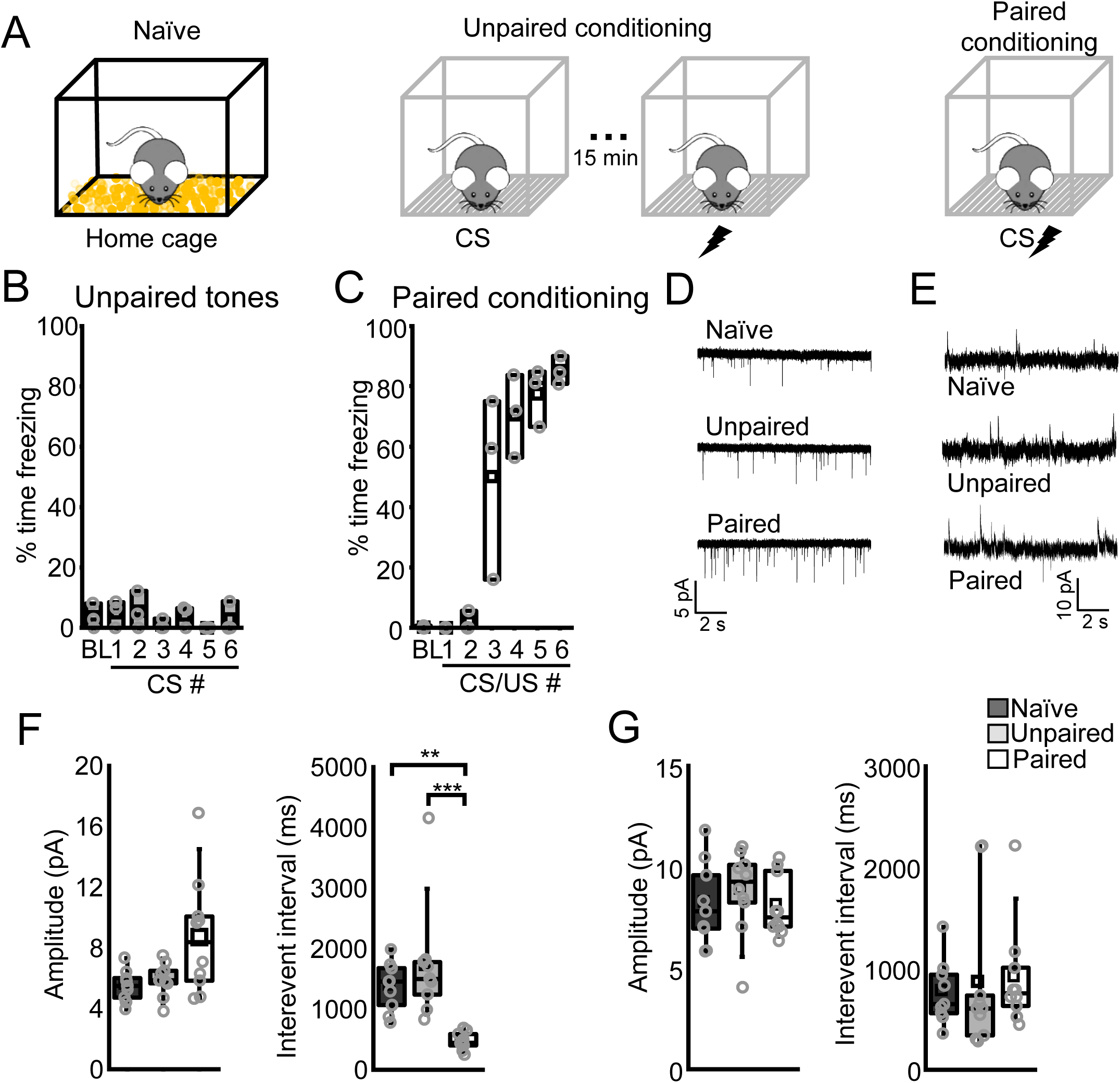
Fear conditioning drives increased glutamatergic transmission onto DP PNs. (A) Mice were either experienced to their home cage (naïve) or subjected to unpaired or paired presentations of a CS (2 kHz, 20 s, 80 dB) and US (2s, 1 mA). Whole-cell recordings were performed 24 hours after training. (D) Representative miniature excitatory postsynaptic current (EPSC) recordings in DP PNs from brain slices prepared from naïve (6 slices from 3 mice), unpaired (5 slices from 3 mice), and paired (6 slices from 3 mice) mice. (E) Representative spontaneous inhibitory postsynaptic current (IPSC) recordings in DP PNs from brain slices prepared from naïve, unpaired, and paired mice. (F) Quantification of the EPSC amplitude and interevent interval for naïve, unpaired, and paired groups. Amplitude: Χ^2^ = 5.44 (2), P = 0.07, Kruskal-Wallis ANOVA, naïve = 10 cells; unpaired = 9 cells; paired = 10 cells. Interevent interval: Χ^2^ = 19.2 (2), P = 6.65 x 10^-5^, Kruskal-Wallis ANOVA; naïve = 10 cells; unpaired = 9 cells; paired = 10 cells. (G) Quantification of the IPSC amplitude and interevent interval for naïve, unpaired, and paired groups. Amplitude: F_2,26_ = 0.46, P = 0.63, one-way ANOVA; naïve = 10 cells; unpaired = 9 cells; paired = 10 cells. Interevent interval: Χ^2^ = 1.77 (2), P = 0.41, Kruskal-Wallis ANOVA; naïve = 10 cells; unpaired = 9 cells; paired = 10 cells. **P < 0.01, ***P < 0.001, Tukey’s or Dunn’s post hoc test. Box plots represent the median (center line), mean (square), quartiles, and 10-90% range (whiskers). Open circles represent data points for individual cells.

**Supplementary Figure 4.**
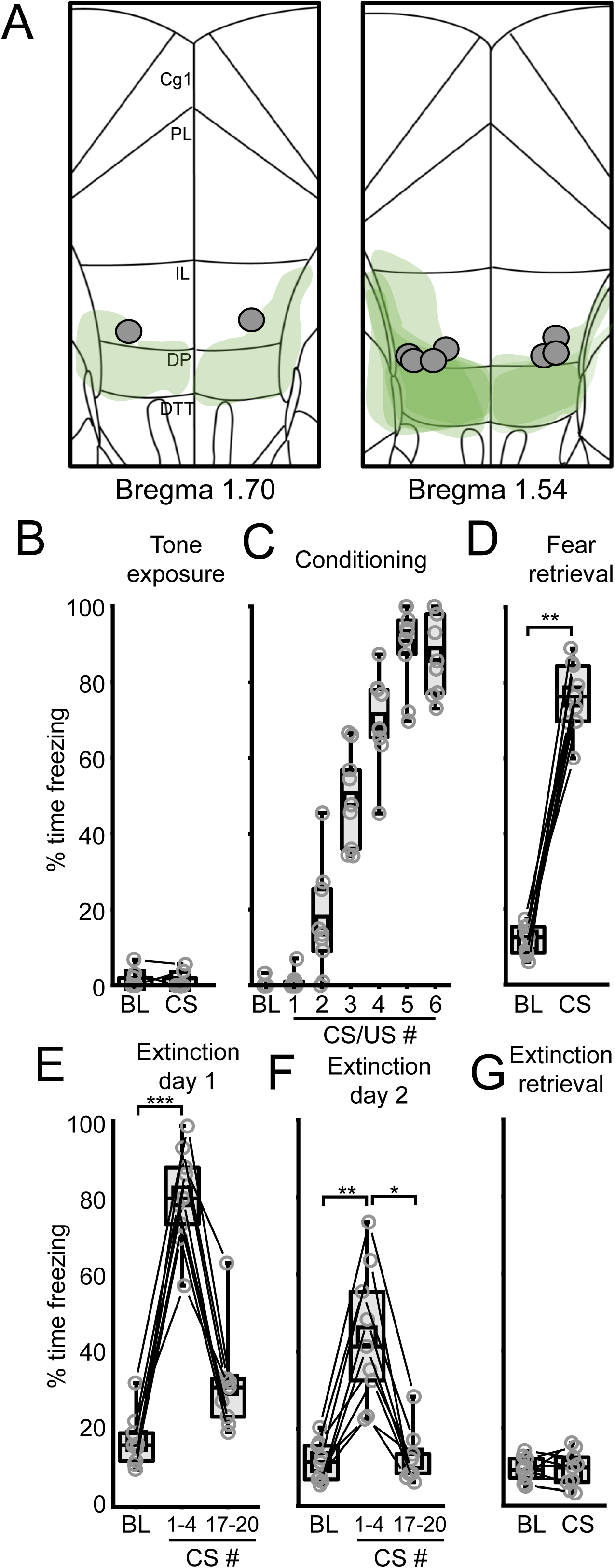
Fiber photometry fiber placements and behavioral data. (A) Fiber placements. Gray circles represent imaging fiber tip termination positions. Transparent green shapes represent extent of viral spread. Quantification of percent time freezing during (B) tone exposure, W = 3, P = 1, Wilcoxen signed rank test, (C) fear conditioning, (D) fear memory retrieval, W = 0, P = 0.004, Wilcoxen signed rank test, (E) extinction day 1, X^2^ (2) = 18, Friedman ANOVA, (F) extinction day 2, X^2^ (2) = 14, Friedman ANOVA, and (G) extinction memory retrieval, t(8) = 0.069, P = 0.95, paired t-test. *, P < 0.05, **, P < 0.01, ***, P < 0.001, Dunn’s post-hoc test. Box plots represent the median (center line), mean (square), quartiles, and 10-90% range (whiskers). Open circles represent data points for individual mice. Lines represent data points from the same animal.

**Supplementary Figure 5.**
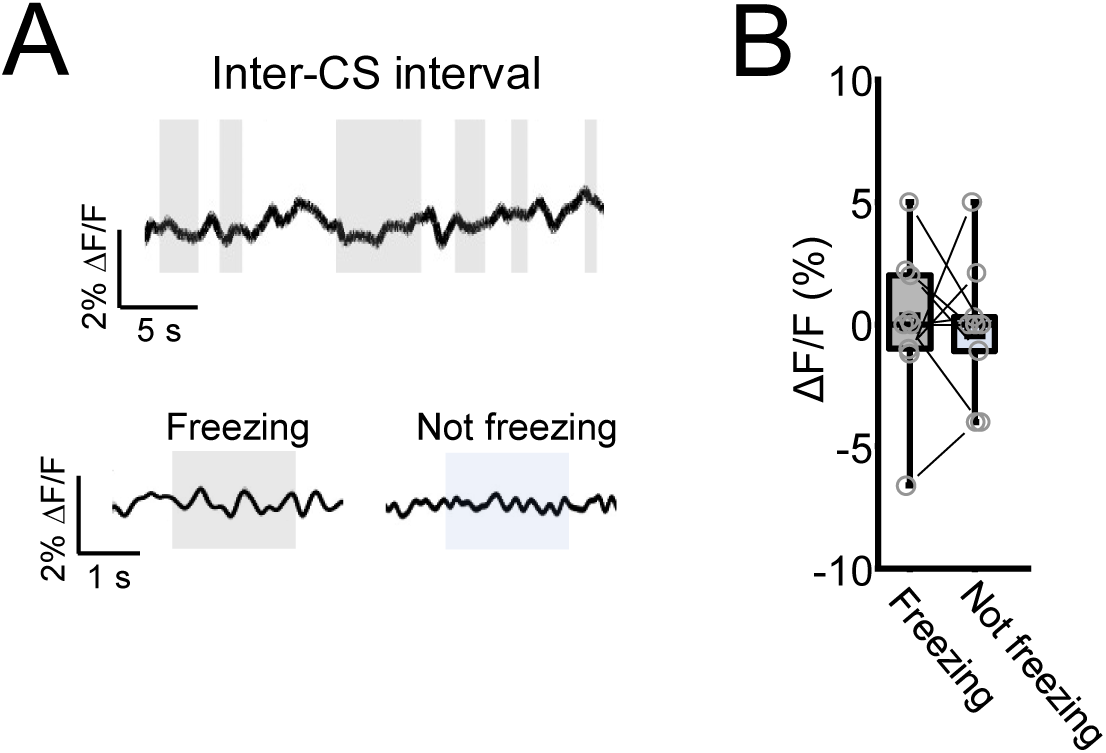
Fiber photometry measurements during spontaneous inter-CS freezing bouts. (A) Top, raw calcium trace during a random 30-s inter-CS interval during the fear memory retrieval test from Figure 3. Gray boxes indicate periods of spontaneous freezing. Bottom, mean aligned calcium traces for epochs where animals were either freezing or not freezing. Gray and blue boxes represent averaged periods where the animal was and was not freezing, respectively. (B) Comparison of ΔF/F percentage between inter-CS epochs where animals were spontaneously freezing or not freezing in the fear memory retrieval test from Figure 3. t(8) = 0.19, P = 0.85, paired t-test. Box plots represent the median (center line), mean (square), quartiles, and 10-90% range (whiskers). Open circles represent data points for individual mice. Lines represent data points from the same animal.

**Supplementary Figure 6.**
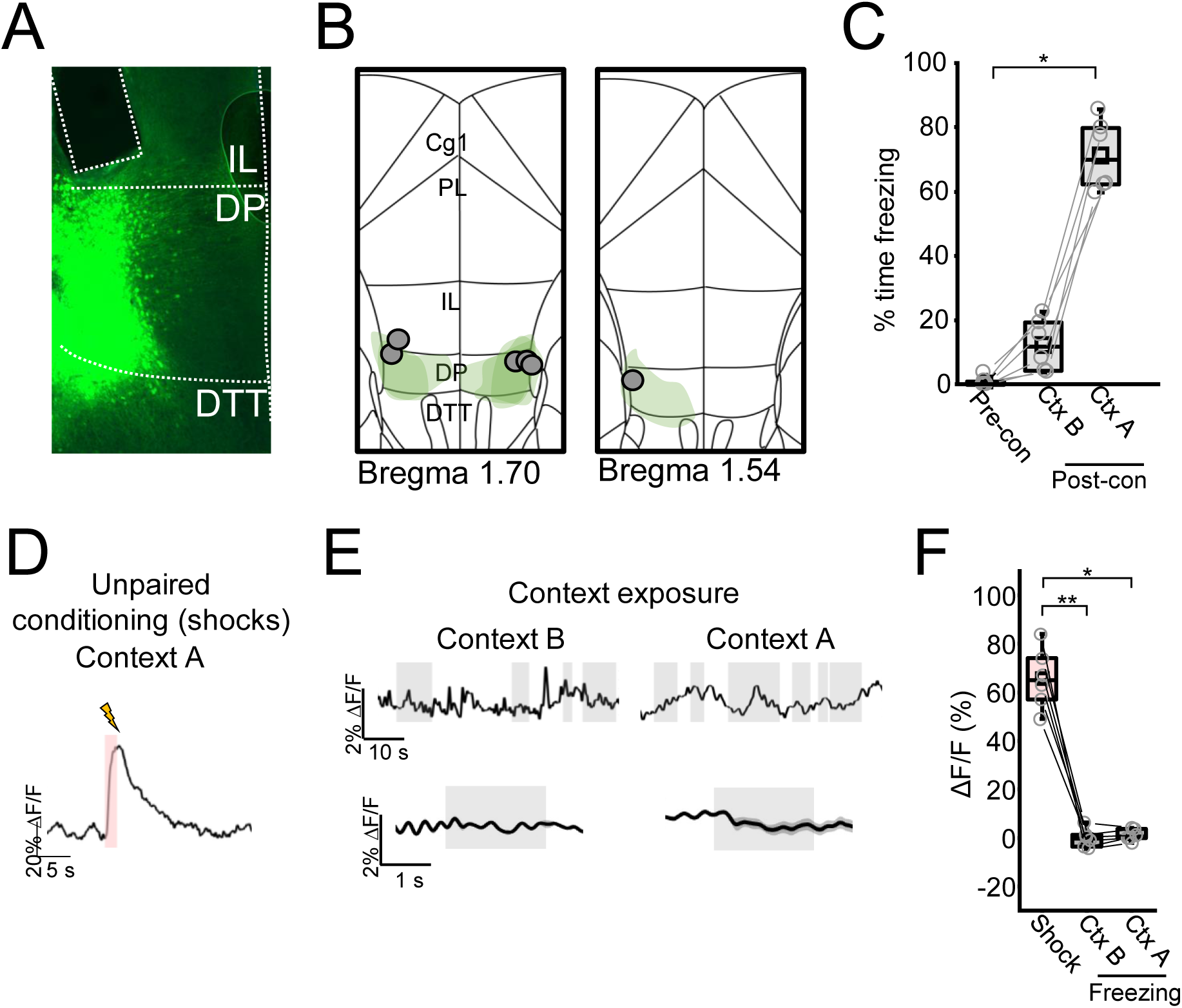
Fiber photometry measurements in DP during unpaired fear conditioning and contextual exposure. (A) Representative histological image of GCaMP8f expression and fiber placement. (C) Mean percent time freezing during pre-conditioning baseline as well as exposure to Context B or Context A at 24 or 48 hours after conditioning, respectively. Effect of context: X^2^ = 12 (2), P = 0.002, Friedman ANOVA. (D) Representative average calcium signal in response to foot shocks from an example animal. Red box denotes foot shock delivery epoch. (E) Top, raw calcium traces during context B or A exposure. Bottom, mean aligned calcium traces for epochs where animals were freezing in each context. Gray boxes indicate periods of freezing. (F) Comparison of calcium signals during shock exposure as well as aligned freezing epochs in contexts A and B. Analysis was restricted to the first two seconds of each epoch normalized to two seconds of the preceding baseline. Effect of shock versus freezing epochs: X^2^ = 10.3 (2), P = 0.006, Friedman ANOVA. *, P < 0.05; **, P < 0.01. Box plots represent the median (center line), mean (square), quartiles, and 10-90% range (whiskers). Open circles represent data points for individual mice. Lines represent data points from the same animal.

**Supplementary Figure 7.**
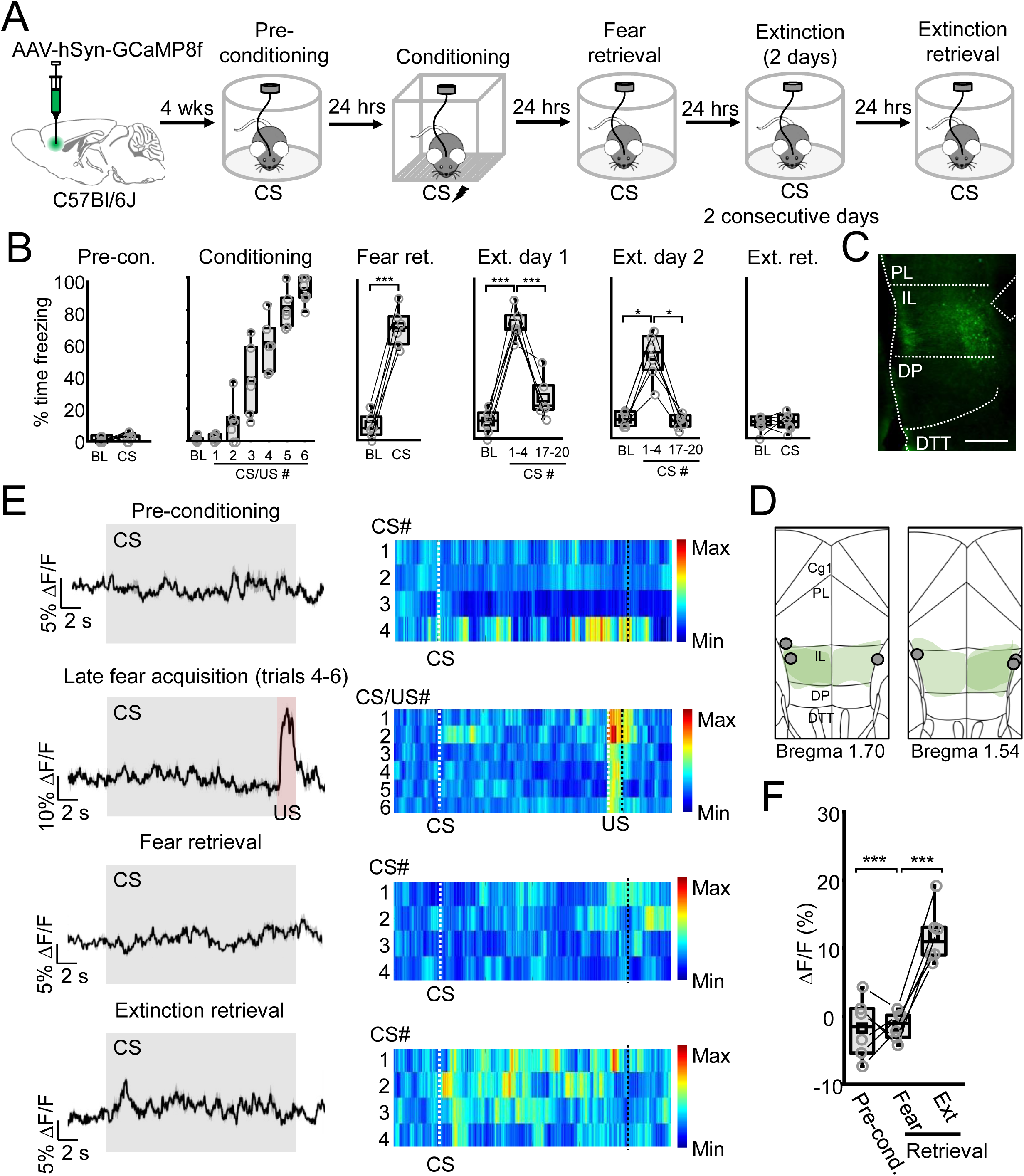
IL signals cue-related activity during extinction memory retrieval. (A) WT mice (n = 6) received infusions of a vector encoding GCaMP8f along with imaging fiber implantation directed at the IL. CS calcium-associated activity was monitored before, during, and after conditioning, in addition to after extinction. Fear conditioning consisted of a baseline period (300 s) followed by 6 pairings of CS (2 kHz, 20 s, 80 dB) and US (2 s, 0.7 mA). Pre-conditioning tone exposure, fear memory retrieval, and extinction memory retrieval all consisted of a baseline period (100 s) followed by four CS presentations (2 kHz, 20 s, 80 dB). (B) Quantification of freezing during pre-conditioning CS exposure (W = 5, P = 0.59), aired fear conditioning, fear memory retrieval (t(5) = -19.3, P = 6.9 x 10^-6^), extinction day 1 (F_2,10_ = 49.5, P = 6.5 x 10^-6^), extinction day 2 (X^2^ (2)= 9, P = 0.01), and extinction memory retrieval (t(5) = -0.32, P = 0.76). (C) Representative histological image of GCaMP8f expression and fiber placement. Scale bar = 400 µm. (D) Schematic representation of GCaMP8f viral expression (green shapes) and fiber placements (gray circles) across all subjects. (E) Left, raw calcium traces averaged across CS trials for each behavior test. Right, heat maps depicting individual calcium responses to each CS trial. (F) Comparison of ΔF/F percentage between pre- and post-conditioning CS presentation, as well as CS-associated activity following extinction training (F_2,10_ = 36.3, P = 2.59 x 10^-5^). *P < 0.05, **P < 0.01, ***P < 0.001, Tukey’s or Dunn’s post hoc test. Box plots represent the median (center line), mean (square), quartiles, and 10-90% range (whiskers). Open circles represent data points for individual mice. Lines represent data points from the same animal.

**Supplementary Figure 8.**
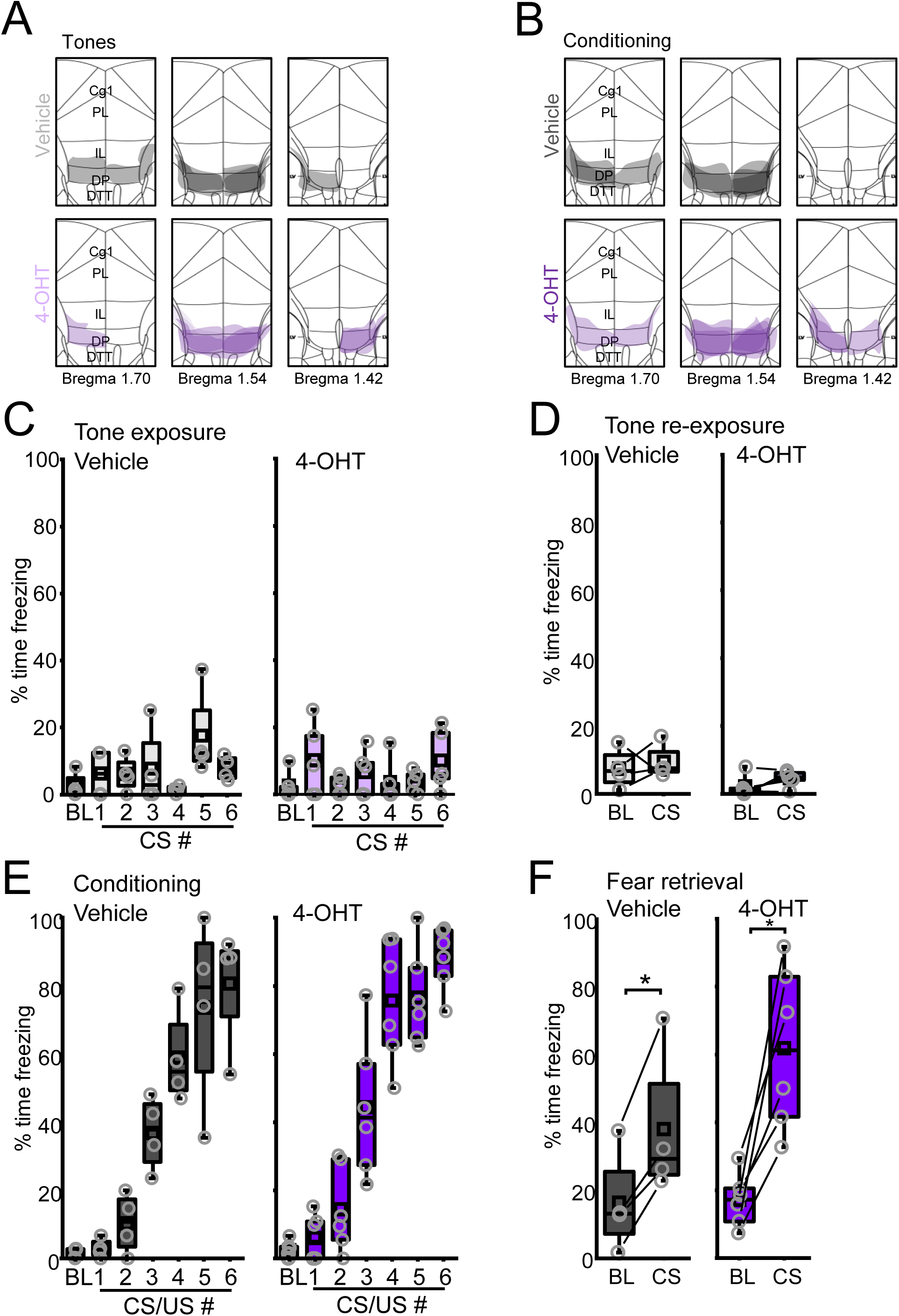
Viral spread and behavior analysis associated with neural tagging data. Representation of viral spread as indicated by mCherry expression for (A) tone-exposed and (B) fear conditioned vehicle- and 4-OHT-injected mice. Quantification of freezing during (C) tone exposure and (D) tone re-exposure (vehicle: t(3) = -0.48, P = 0.67, paired t-test; 4-OHT: W = 3, P = 0.31, Wilcoxen signed rank test). Quantification of freezing during (E) fear conditioning and (F) fear memory retrieval (vehicle: t(3) = -5.23, P = 0.014, paired t-test; 4-OHT: W = 0, P = 0.03, Wilcoxen signed rank test). Box plots represent the median (center line), mean (square), quartiles, and 10-90% range (whiskers). Open circles represent data points for individual mice.

**Supplementary Figure 9.**
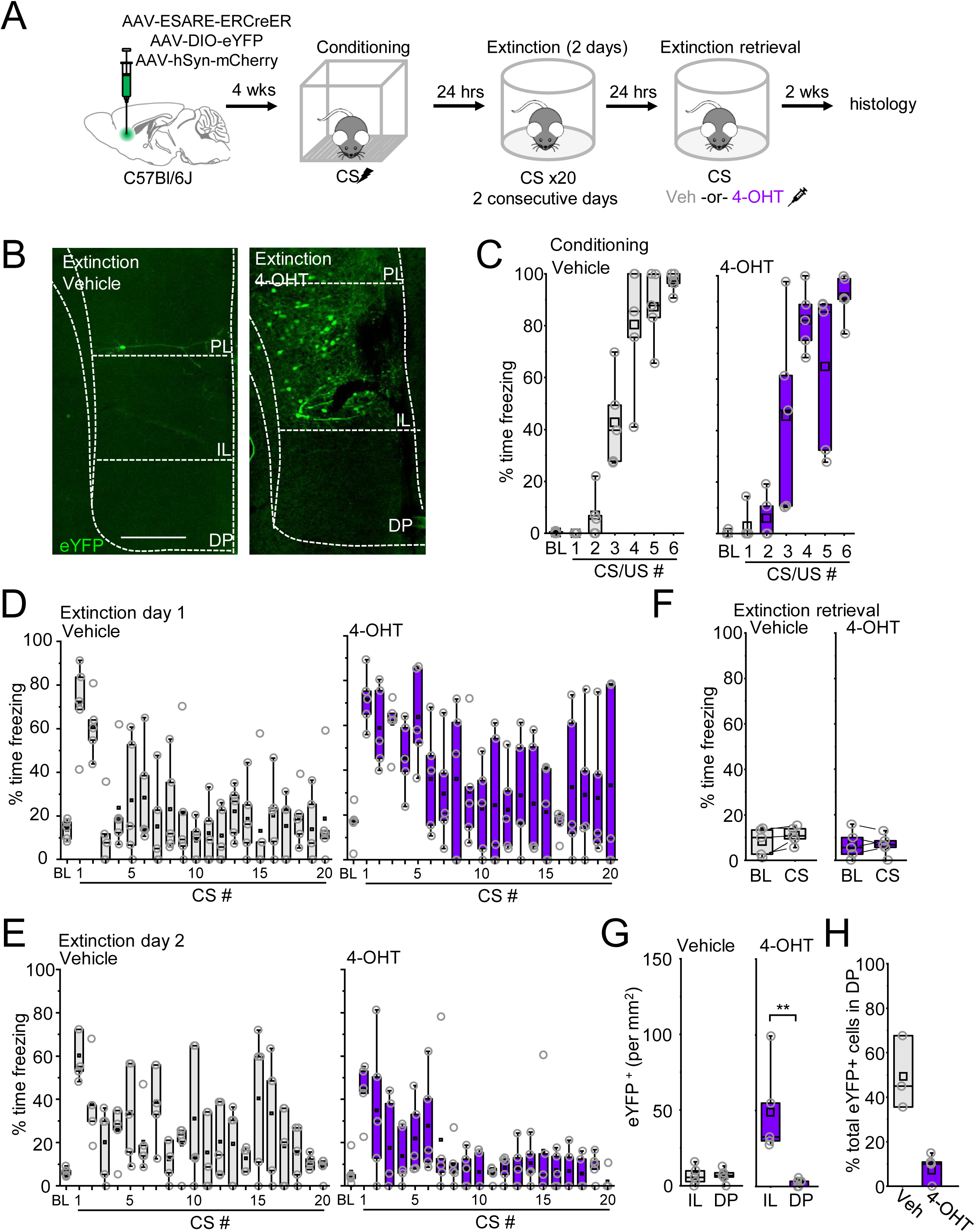
Behavior analysis and histology for neural tagging in response to extinction training. (A) Behavioral paradigm for tagging infralimbic (IL) and dorsal peduncular (DP) cortex neurons following extinction training. (B) Representative histological images for animals subjected to extinction training and injected with either vehicle or 4-hydroxytamoxifen (4-OHT). Scale bar = 500 µm. Quantification of freezing during (C) fear conditioning, (D) day 1 of extinction training, (E) day 2 of extinction training, and (F) extinction memory retrieval (vehicle: t(4) = -1.81, P = 0.14, paired t-test; 4-OHT: t(4) = 0.02, P = 0.99, paired t-test). (G) Quantification of the number eYFP+ neurons in the IL and DP (vehicle: t(8) = 0.008, P = 0.99, two-sample t-test; 4-OHT: U = 25, P = 0.012, Mann-Whitney U-test). (H) Quantification of the fraction of total eYFP+ neurons that are localized in the DP (t(6) = -5.24, P = 0.002, two-sample t-test). Box plots represent the median (center line), mean (square), quartiles, and 10-90% range (whiskers). Open circles represent data points for individual mice.

**Supplementary Figure 10.**
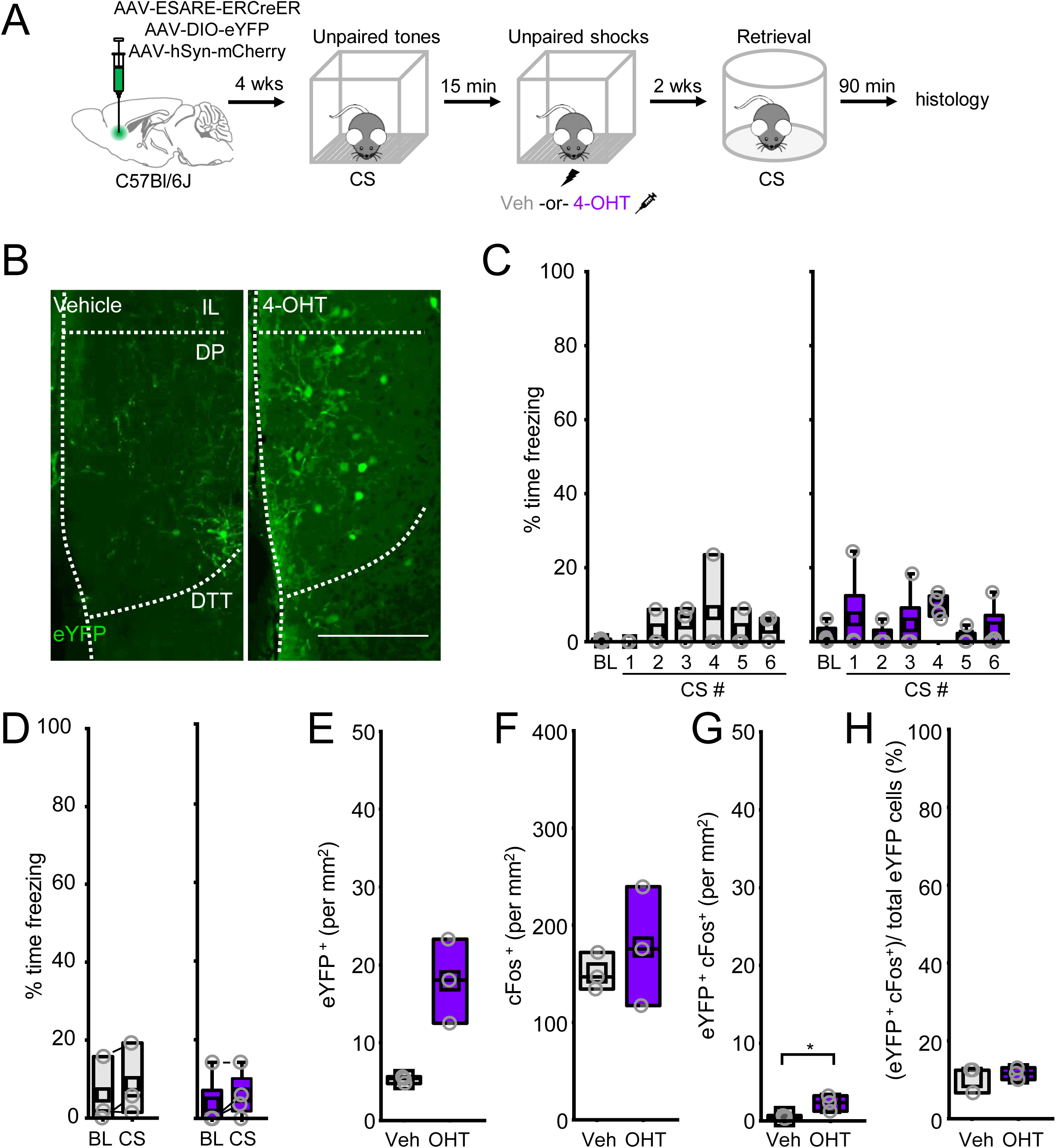
Behavior analysis and histology for neural tagging in response to unpaired conditioning. (A) Four weeks after vector infusion, mice were subjected to unpaired fear conditioning and injected with vehicle or 4-OHT immediately after shock delivery. Following 2 weeks, mice were subjected to a CS-evoked fear memory retrieval test and tissue was processed for cFos staining 90 minutes later. (B) Representative histological images from vehicle- and 4-OHT-injected mice. Scale bar = 500 µm. (C) Quantification of the percent time freezing during unpaired CS presentation for vehicle (gray) and 4-OHT (purple) mice. (D) Quantification of the percent time freezing during CS-evoked fear memory test for vehicle (t(2) = -1.63, P = 0.25, paired t-test) and 4-OHT (W = 0, P = 0.25, Wilcoxen signed-rank test). Comparisons between unpaired vehicle (n = 4 mice) and unpaired 4-OHT mice (n = 3) for (E) eYFP+ (U = 0, P = 0.081, Mann-Whitney U test), (F) cFos+ (t(4) = -0.71, P = 0.52), (G) eYFP+ and cFos+ (t(4) = -2.96, P = 0.041), and (H) eYFP+ and cFos+/ eYFP+ (U = 4, P = 1, Mann-Whitney U-test). Box plots represent the median (center line), mean (square), quartiles, and 10-90% range (whiskers). Open circles represent data points for individual mice.

**Supplementary Figure 11.**
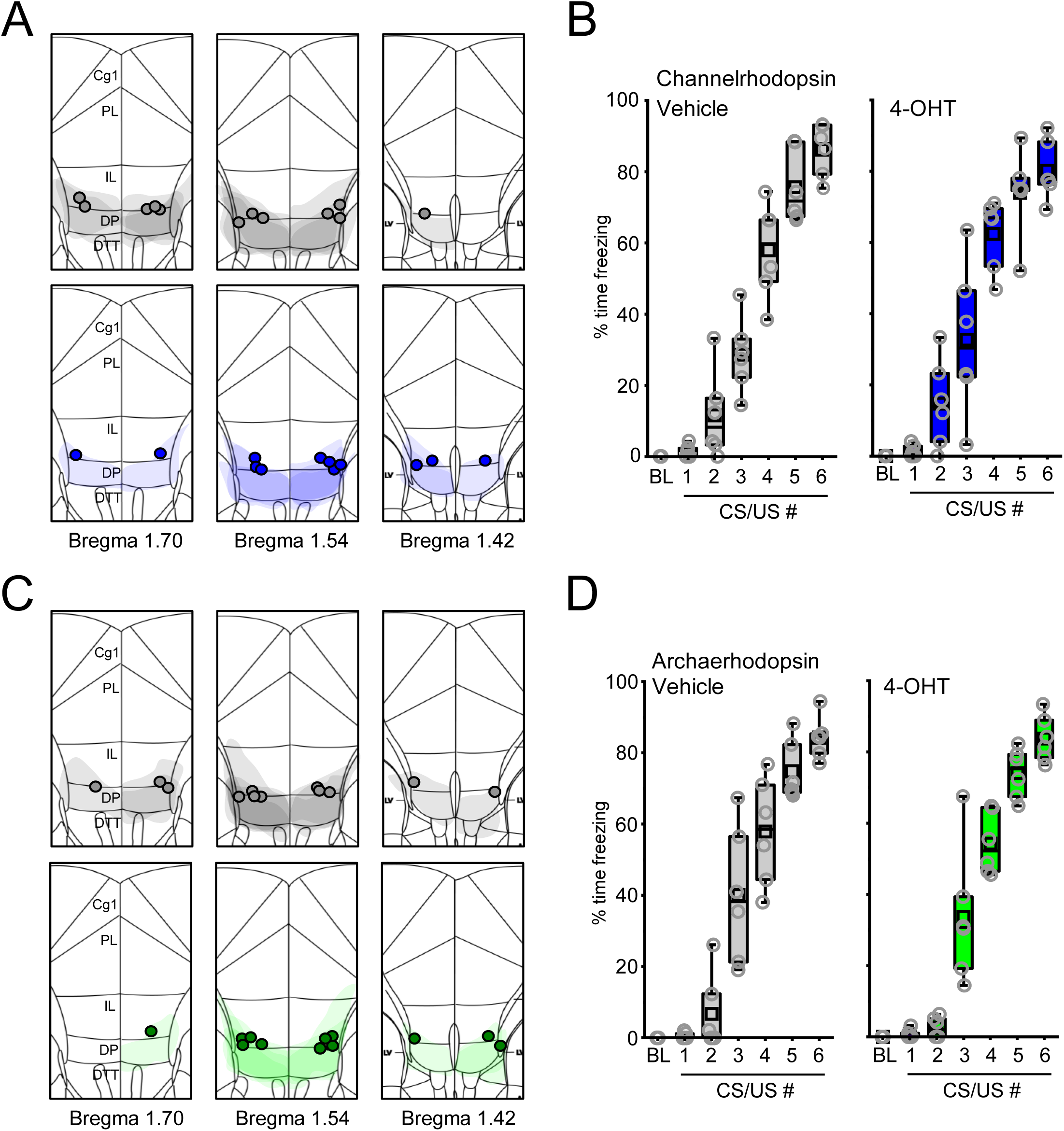
Fiber placements, viral expression, and behavioral data for optogenetic experiments. (A) Optic fiber placements and viral expression for channelrhodopsin-expressing mice. Gray and blue circles and shapes represent optic fiber tip termination positions and viral expression patterns as indicated by mCherry expression for vehicle- and 4-OHT-injected mice, respectively. (B) Quantification of percent time freezing during fear conditioning for vehicle- (gray) and 4-OHT- (blue) injected channelrhodopsin-expressing mice. (C) Optic fiber placements and viral expression for archaerhodopsin-expressing mice. Gray and green circles represent optic fiber tip termination positions and viral expression patterns as indicated by mCherry expression for vehicle- and 4-OHT-injected mice, respectively. (D) Quantification of percent time freezing during fear conditioning for vehicle- (gray) and 4-OHT- (green) injected archaerhodopsin-expressing mice. Box plots represent the median (center line), mean (square), quartiles, and 10-90% range (whiskers). Open circles represent data points for individual mice.

**Supplementary Figure 12.**
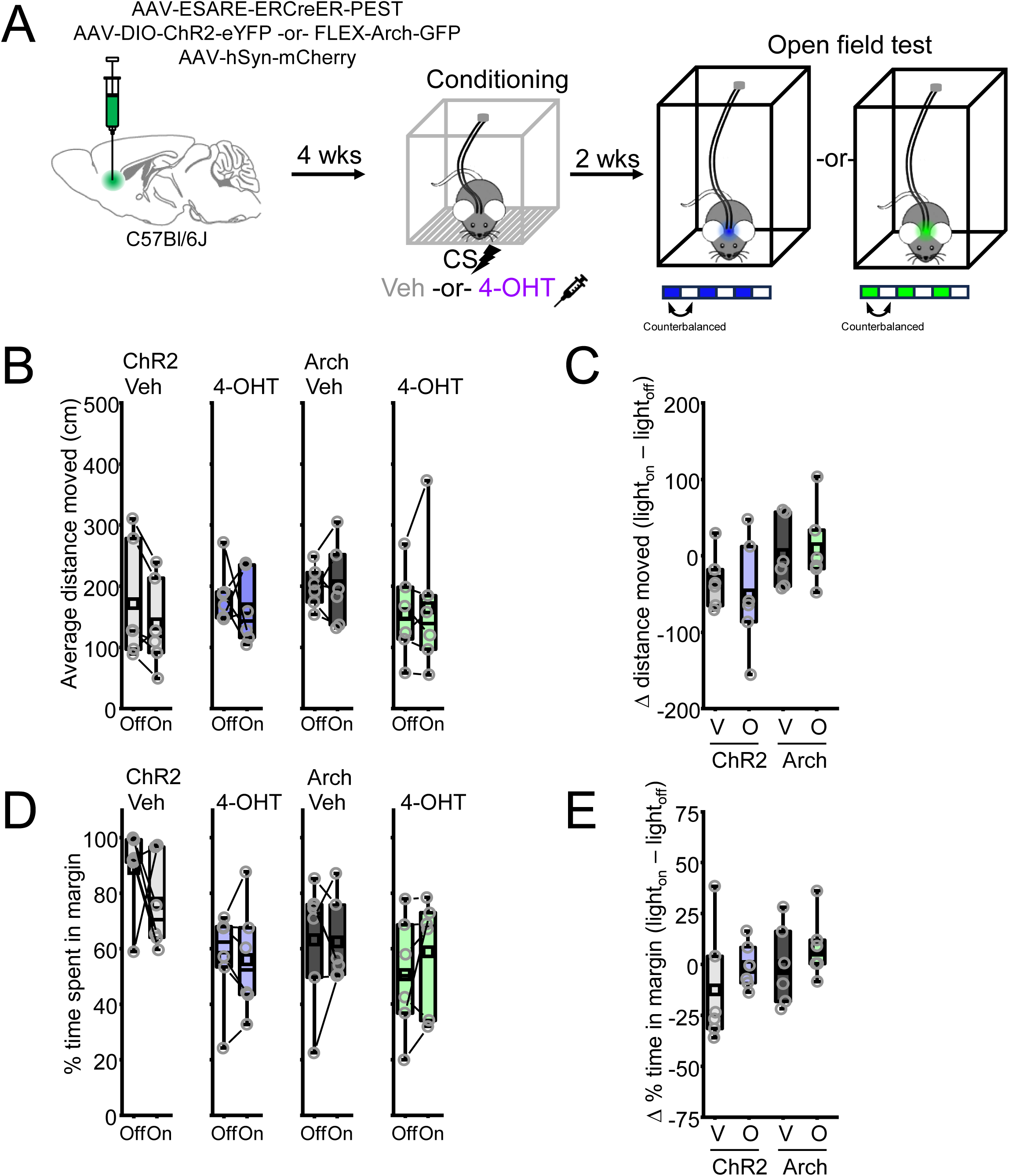
Optogenetic manipulations during the open field test. (A) Neural tagging and manipulation paradigm. Four weeks after vector infusion and optic fiber implantation, mice were subjected to paired fear conditioning and injected with vehicle or 4-OHT immediately after. Following 2 weeks, mice were placed in an open field for 10 minutes and optogenetic activation (for ChR2-injected mice: 473 nm, 20 Hz, 5 ms pulses, 1 min on/off epochs) or silencing (for Arch-injected mice: 532 nm, solid light, 1 min on/off epochs) was performed. (B) Quantification of distance moved during light on and light off epochs for vehicle- (t(5) = 2.23, P = 0.076, paired t-test) and 4-OHT- (t(5) = 0.62, P = 0.56, paired t-test) injected ChR2 mice and vehicle- (t(5) = -0.09, P = 0.93, paired t-test) and 4-OHT- (t(5) = -0.43, P = 0.69, paired t-test). (C) Quantification of the change in distance moved was performed by subtracting distance during light_on from light_off (F(3) = 1.71, P = 0.2). V is vehicle, O is 4-OHT. (D) Quantification of the percent time spent in the open field margin during light on and light off epochs for vehicle- (W = 14, P = 0.56, Wilcoxen signed-rank test) and 4-OHT- (t(5) = 0.2, P = 0.85, paired t-test) injected ChR2 mice and vehicle- (t(5) = 0.09, P = 0.93, paired t-test) and 4-OHT- (t(5) = - 1.32, P = 0.24, paired t-test). (E) Quantification of the change in the percent time spent in the margin was performed by subtracting percent time in margin during light_on from light_off (F(3) = 1.11, P = 0.37, one-way ANOVA). V is vehicle, O is 4-OHT. Box plots represent the median (center line), mean (square), quartiles, and 10-90% range (whiskers). Open circles represent data points for individual mice.

**Supplementary Figure 13.**
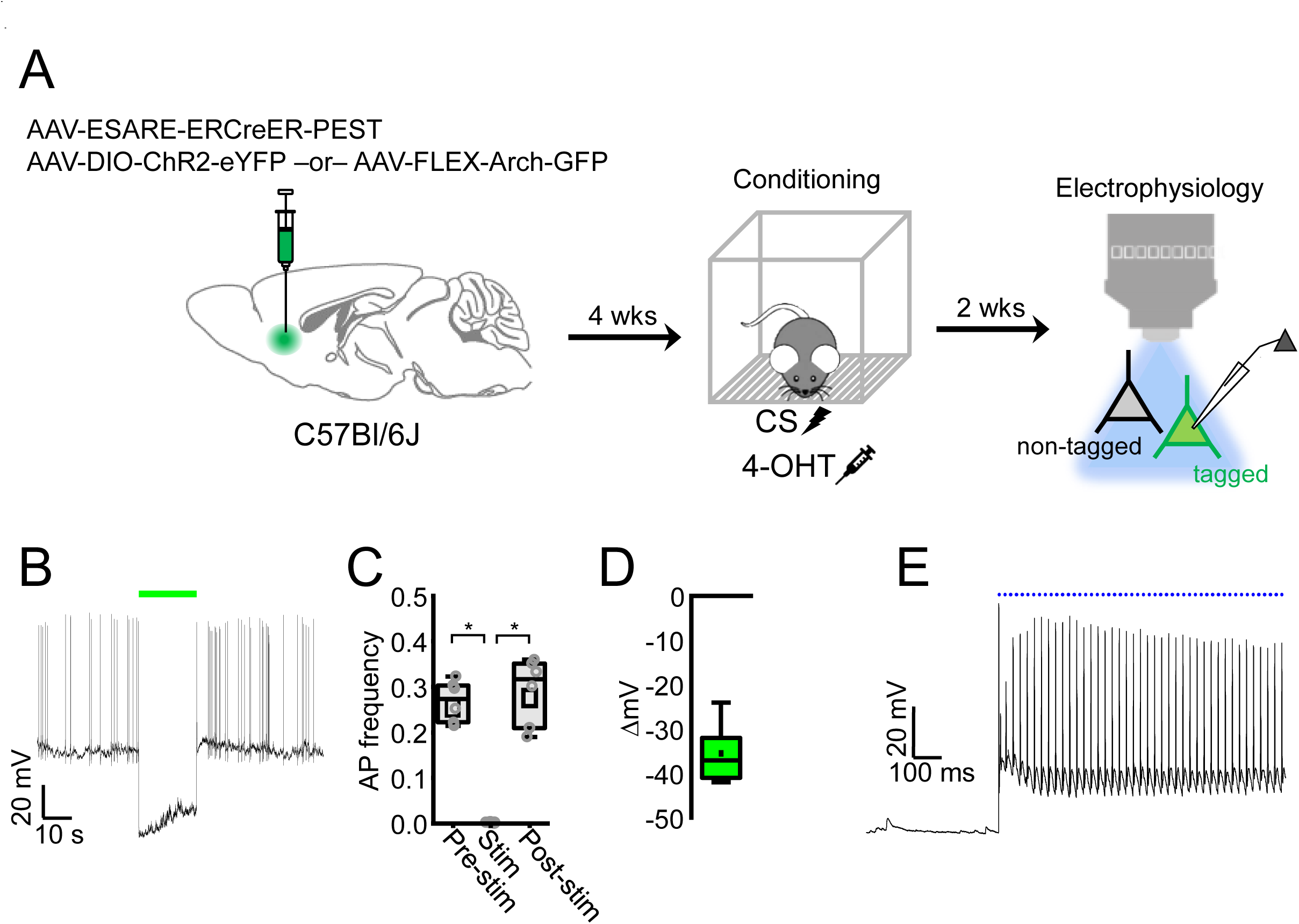
Electrophysiological validation of optogenetic manipulations. (A) ESARE-ERCreER and a Cre-dependent ChR2-eYFP or Cre-dependent Arch-GFP was infused into the DP of wild-type male and female mice. Mice were subjected to paired fear conditioning 4 weeks later and immediately injected with 4-OHT after training. After 2 weeks, acute brain slices were prepared and recordings were made from ChR2- or Arch-expressing fear-tagged DP neurons. (B) Example recording of spontaneous action potential firing from a neuron expressing Arch with and without optogenetic silencing (532 nm; solid light, 20 s epoch; denoted with green line). (C) Quantification of spontaneous action potential firing before, during, and after optogenetic silencing. X^2^ (2) = 0.009, Friedman ANOVA. n = 6 cells from 3 mice. (D) Quantification of the change in steady-state membrane potential during optogenetic silencing compared to a pre-stim baseline. n = 6 cells from 3 mice. (E) Representative recording of action potentials elicited by pulses of blue light (473 nm; 20 Hz, 5 ms pulses, 20 s epochs) in ChR2-expressing tagged neurons. n = 8 cells from 3 mice. *, P < 0.05, Dunn’s post-hoc test. Box plots represent the median (center line), mean (square), quartiles, and 10-90% range (whiskers). Open circles represent data points for individual cells.

